# Genome wide transcriptional changes underlie gradual and recurrent adaptation to protein malnutrition in zebrafish

**DOI:** 10.64898/2026.02.26.708142

**Authors:** Siyao Wang, Laura Childers, Fernando Martinez, Michel Bagnat, Jieun Park

**Affiliations:** Department of Cell Biology, Duke University, Durham, NC 27710, USA; Department of Molecular Genetics and Microbiology, Duke University, Durham, NC 27710, USA; Neuroscience Center, University of North Carolina, Chapel Hill, NC, 27599, USA; Department of Genetics, University of North Carolia at Chapel Hill, Chapel Hill, NC, 27599, USA

## Abstract

Lysosome-rich enterocytes (LREs) are a specialized population of intestinal cells that mediate the uptake and absorption of dietary proteins in fish and neonatal mammals. Loss of LRE function causes inhibited growth and poor survival due to protein malnutrition. Previously, we reported that in zebrafish loss of Plasmolipin (pllp), an endosomal membrane protein highly expressed in LREs, impairs LRE differentiation and dietary protein absorption, resulting in marked reduction in survival rates. Raising *pllp* homozygous mutants surviving to adulthood and in-crossing them for multiple generations resulted in their adaptation to malnutrition, with adapted *pllp* mutants showing no survival deficits. To uncover mechanisms underlying this phenomenon, we compared the older adapted *pllp* allele with genetically related wild type (WT) fish and a newly generated *pllp* mutant allele. Using transcriptome profiling and quantitative protein absorption assays, we found that adapted *pllp* mutants exhibit upregulation of LRE endocytic components, resulting in a capacity for protein absorption that exceeds that of WT. This hyperactivation of LRE endocytic and absorptive activity is aided by a fine-tuned transcriptional regulation of immune genes that may contribute to the enhanced survival of *pllp* mutants in the face of increased exposure to environmental antigens. Genetic analyses indicate that these adaptations emerge gradually and are recurrent as shown experimentally by the adaptation of a mutant allele upon inbreeding and natural selection. Overall, our study illustrates that genome wide transcriptional changes underly adaptation mechanisms that enhance intestinal function and organismal survival in response to protein malnutrition.

**Author Summary:** In this study, we investigated how zebrafish adapt to protein malnutrition when the function of specialized intestinal protein-absorbing enterocytes, also found in newborn mammals, is impaired. We observed that fish carrying a mutation that severely disrupts intestinal protein absorption gradually recovered their ability to survive over multiple generations of inbreeding, even though the underlying mutation remained intact. By comparing gene activity in these adapted fish with that of newly generated mutants, we found that adaptation involves a coordinated rewiring of two systems: the enterocytes themselves became hyperactivated, absorbing more protein than even wildtype fish, while the immune system was simultaneously recalibrated to dampen inflammation. We further showed that this adaptive process is recurrent by using a second, independently generated mutant line that underwent a strikingly similar recovery trajectory over successive generations. Together, our findings reveal that animals can overcome a severe, heritable nutritional deficit through a gradual, genome-wide transcriptional response that fundamentally reshapes intestinal function across generations.

## Introduction

As an essential macronutrient, dietary protein is critical for animal development and for sustaining normal organismal physiology. Protein intake deficiencies, clinically manifested as stunting, wasting, and severe syndromes such as Kwashiorkor (nutritional edema), remain a major global health concern that contribute to pediatric morbidity and mortality (1,2). Dietary protein absorption in zebrafish and neonatal mammals is highly dependent on lysosome-rich enterocytes (LREs), which are a group of specialized epithelial cells located in the ileum that are characterized by their giant lysosomal vacuoles and highly endocytic activity and capacity (3). Previously described as vacuolated or neonatal enterocytes (4–6), LREs are widely conserved across various species of mammals including humans, rat, sheep, and pigs, as well as in teleost fishes(6–10). While LREs are replaced by regular adult-type enterocytes upon weaning in mammals, they are retained through adulthood in fishes, which makes zebrafish an ideal model to study these cells that are only transient in the neonatal mammalian gut (3,11).

LREs absorb dietary protein macromolecules via a multi-ligand complex machinery consisting of the endocytic adaptor protein Dab2, the transmembrane linker protein amnionless (Amn), and the scavenger receptor cubulin (Cubn)(3). Loss of these components leads to poor survival, stunted growth in zebrafish and mice and also to intestinal edema reminiscent of Kwashiorkor disease in mice (3). Apart from the endocytic machinery, the correct functioning of LREs also relies on endosomal protein plasmolipin (Pllp) (12). Pllp is enriched at the apical region of epithelial <u>Madin-Darby Canine Kidney</u> (MDCK) cells during polarization in 3D cultures, where it is critical for maintaining apical sorting, recycling of SNAREs, and the activation of Notch signaling pathway (12). In zebrafish, disruption of Pllp function results in impaired endocytic activity in LREs, leading to markedly reduced survival during larval stages (12).

Adaptation to nutrient scarcity occurs across two distinct timescales: the acute physiological response within a single individual and the transgenerational adaptation that shapes the fitness of subsequent generations. Severe acute malnutrition (SAM) and protein-energy malnutrition (PEM) in children and neonatal animal models are characterized by a shift to “protein-sparing” energy use, reduced growth, intestinal epithelial remodeling, endocrine changes, fatty liver, hypometabolism, and higher mortality during infections, which are often interpreted as adaptive attempts to survive at the cost of growth and immune competence(13–16). At the core of adaptation are several well-studied nutrient-sensing and autophagy pathways including mammalian target of rapamycin (mTOR), AMP-activated protein kinase (AMPK), and General Control Non-depressible 2 (GCN2) (17–20). Interestingly, in SAM or PEM animal models, refeeding alone often fails to return the animals to normal growth or cellular functions (13, 14), suggesting that the acute episode leaves a durable physiological imprint on individuals. Previous studies demonstrated that such episodes remodel chromatin at autophagy and metabolic genes via DNA methylation and histone modifications, generating an epigenetic memory that is partially heritable(21,22). Indeed, in murine models, offsprings produced by parents fed with low-protein diet (LPD) or with restricted access to dietary protein were shown to exhibit changes in promoter methylation and altered expressions of genes associated with metabolism, immune regulations, oxidative stress and neurodevelopment, with effects often strongest in F1 and decreasing to near normalization by F3(23–26). Most transgenerational studies on protein deficiency focus on manipulating wildtype animals using formulated low protein diets. In those studies, the transgenerational adaptation effects are largely attenuated by the F3 (23,26–30). It was unclear how animals with inheritable genetic deficits that impair dietary protein intake could overcome a lethal survival disadvantage through lasting transgenerational adaptation that fundamentally changes the organ’s functional capacity.

Serendipitously, we found that raising surviving homozygous *pllp* mutant zebrafish and subsequent inbreeding led to a loss of the survival defect originally reported in these mutants. Here, we investigate how these adapted *pllp* mutants could recover from a strong and largely lethal nutritional deficit via multigenerational adaptation and selection. We show that the F5 and subsequent generations of this zebrafish line show persistent changes in the transcriptional profiles of genes involved in protein absorption and immune responses that are in sharp contrast to that of newly generated *pllp* mutant alleles. Furthermore, using quantitative protein uptake and degradation assays, we show that the LREs in these adapted fish exhibit surprising hyperactivation of their protein endocytosis and degradation capabilities. Moreover, genetic experiments revealed that the adaptation to protein malnutrition is recurrent. Together, our results reveal that concurrent transcriptional regulations of multiple genes spanning LRE specific and as well as systemic processes underly transgenerational adaptation to protein malnutrition in zebrafish.

## Results

### Zebrafish with impaired survival and reduced intestinal function due to the loss of *pllp* improve under multigenerational selection pressure

Previously, in Rodríguez-Fraticelli et al. (2015) we reported that zebrafish with a mutation in *pllp* (expression shown in Figure 1A), named *pllp^pd1116^-/-,* showed severely impaired survival performance as well as reduced endocytosis of fluorescently labeled dextran in LREs. After creation of the original *pd1116*-/- stock, this genetic line was maintained via in-crossing of the surviving adult homozygous mutants. Following its generation, the in-cross offspring of F1 *pd1116*-/- was tested again for its survival performance using newly defined custom diets (Figure 1B) (31). Six days-post-fertilization (6 dpf) larvae were fed with either a calorie-restricted (10 mg food/tank/day) high-protein (HP) diet or calorie-restricted low-protein (LP) diet for 10 days until 16 dpf. Although the compromised survival phenotype originally reported in our previous study persisted when compared to the parent wild-type (WT) EK strain, the survival rate of those F2 *pd1116*-/- was noticeable higher than what we originally documented using undefined diets (Figure 1C). Subsequently, the mutant strain was maintained as a homozygous mutant for 4 additional generations (Figure 1B). Then, to use them as reference in a new study, experimental in-crosses were generated and tested for survival using our defined diets and following the same calorie restriction scheme used previously. Surprisingly, the survival rate of F6 *pd1116*-/- offspring was comparable to EK under either diet condition (Figure 1D). Upon sequencing the *pllp* locus, we found no changes or loss to the original *pd1116*-/- mutation site compared to previously published data (Figure S1A, B).

**Figure 1.**
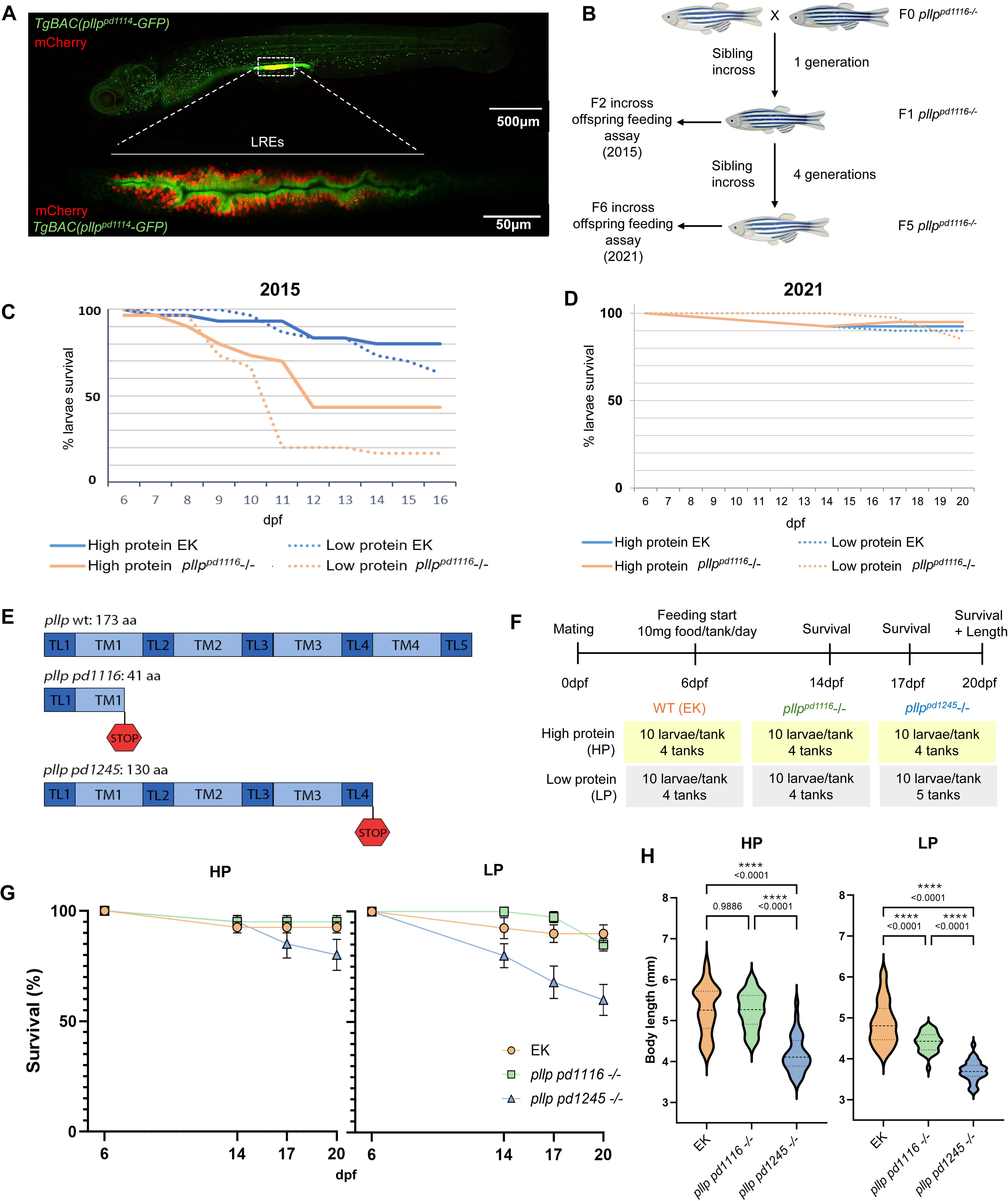
Increased survival of *pd1116*-/- under generational selection pressure. **(A)** Live image of a *TgBAC(pllp^pd1114^-GFP)* 6dpf larva gavaged with mCherry protein. The magnified area below is the LRE region. **(B)** Pedigree of the *pd1116* allele showing the generations used for feeding assays. **(C)** Survival curve of *pd1116*-/- vs wild-type (EK) fed with calorie-restricted (10mg/tank/day) high-protein diet and calorie-restricted low-protein diet; experiment conducted in 2015; n=4 tanks/condition/genotype, starting 10 larvae/tank. **(D)** Survival curve of *pd1116*-/- vs EK conducted in 2021; n=4 tanks/condition/genotype, starting 10 fish/tank. **(E)** Diagram of *pllp* alleles; *pd1116* allele was created by TALENs and has an early stop at amino aa41; *pd1245* was created by CRISPR-Cas9 using a single gRNA and has an early stop at aa130. TL=topological domain; TM=transmembrane domain. **(F)** Schematics of feeding experiment with calorie-restricted high-protein diet (HP) and calorie-restricted low-protein (LP) diet. **(G)** Survival curves of 3 genotypes over 20 days under HP or LP diet. n=4 tanks/condition/genotype except for *pd1245*-/- LP (n=5 tanks), starting 10 fish/tank. **(H)** Body lengths of surviving fish of 3 genotypes at 20dpf. n=37 for EK HP; n=38 for *pd1116*-/- HP; n=32 for *pd1245*-/- HP; n=36 for EK LP; n=34 for *pd1116*-/- LP; n=30 for *pd1245*-/- LP. One-way ANOVA.

To investigate the mechanisms underlying *pd1116*-/-’s survival adaptation, we established a new loss of function (LOF) allele, *pd1245,* to mimic *pd1116*-/- in its pre-adapted state. Given the lack of suitable CRISPR target sites in exon1, which harbors the original *pd1116* TALEN-generated mutation, *pd1245* was created using a single CRISPR gRNA targeting exon 2 of *pllp*, resulting in a 10 base pair (bp) deletion which produces a nonsense mutation downstream at amino acid 130 and a truncation of transmembrane domain 4 (Figure 1E). To characterize the new *pd1245* allele, we first performed RT-PCR to examine transcript expression levels and found that both *pd1245* and *pd1116* alleles exhibit significant degradation of the *pllp* transcript as compared to WT EK, suggesting that non-sense mediated decay (NMD) occurs as a result of the presence of premature stop codons (Figure S1C, C’). Integrative Genomics Viewer (IGV) visualization of splice junction coverage combined with Differential Exon Usage Sequence (DEXseq) analysis of relative exon usage at the *pllp* locus revealed the absence of novel splice isoforms or exon skipping, excluding restored *pllp* functioning by skipped-over mutation site or alternative splicing as the underlying cause of *pd1116*-/-’s adaptation (Figure S1D-F).

Next, we conducted a 20-day feeding experiment using a calorie-restricted HP or LP diet design. At 6 dpf, larvae from in-cross matings of EK, *pd1116*-/- or *pd1245*-/- were sorted into 4 or 5 tanks per genotype per diet condition at a density of 10 larvae per tank. Each tank received 10 mg of HP or LP diet per day until 20 dpf. The number of surviving larvae per tank were counted at 14, 17, and 20 dpf to record survival performance. Standard length (32) of any remaining larvae at 20 dpf was measured as proxy of growth (Figure 1F). We found that the average survival rate of *pd1245*-/- larvae declined to 80% under HP and 60% under LP, as compared to 92.5% (HP) and 90% (LP) for EK or 95% (HP) and 85% (LP) for *pd1116*-/- (Figure 1G). Furthermore, body growth of *pd1245*-/- larvae was significantly stunted (4.18mm HP, 3.70mm LP) as compared to both EK (5.26mm HP, 4.88mm LP) or *pd1116*-/- (5.25mm HP, 4.41mm LP) under either feeding condition (Figure 1H). Together, these data show that *pd1245*-/- partially reproduced the impaired survival phenotype and could be used as an approximation of the *pd1116*-/- in its pre-adapted state comparative studies.

### Transcriptomic profiling reveals genotype-dependent and diet-dependent gene expression programs in *pllp* alleles

To identify transcriptomic changes associated with *pllp* loss and adaptation, we performed RNA sequencing on 20 dpf whole zebrafish larvae from three genotypes (EK, adapted *pd1116*-/-, and newly generated *pd1245*-/-) and feeding conditions described in Figure 1F. Hierarchical clustering of all expressed genes revealed distinct transcriptional profiles across genotype and diet conditions (Figure 2A). Principal component analysis (PCA) of global gene expression patterns showed that PC1 separated samples primarily by genotype, with *pd1116*-/- clustering closer to EK than to *pd1245*-/-, consistent with transcriptional normalization occurring in the adapted *pd1116*-/- strain (Figure 2B). Notably, PC1 also captured diet-dependent separation within mutant genotypes as both *pd1116*-/- and *pd1245*-/- showed clear separation between HP and LP conditions, whereas EK HP and LP samples clustered together. This indicates that while diet has minimal impact on WT transcription, it profoundly influences gene expression in both *pllp* mutant backgrounds. PC2 further resolved genotype-by-diet interactions, separating *pd1116*-/- and *pd1245*-/- into distinct transcriptional states under each dietary condition (Figure 2B).

**Figure 2.**
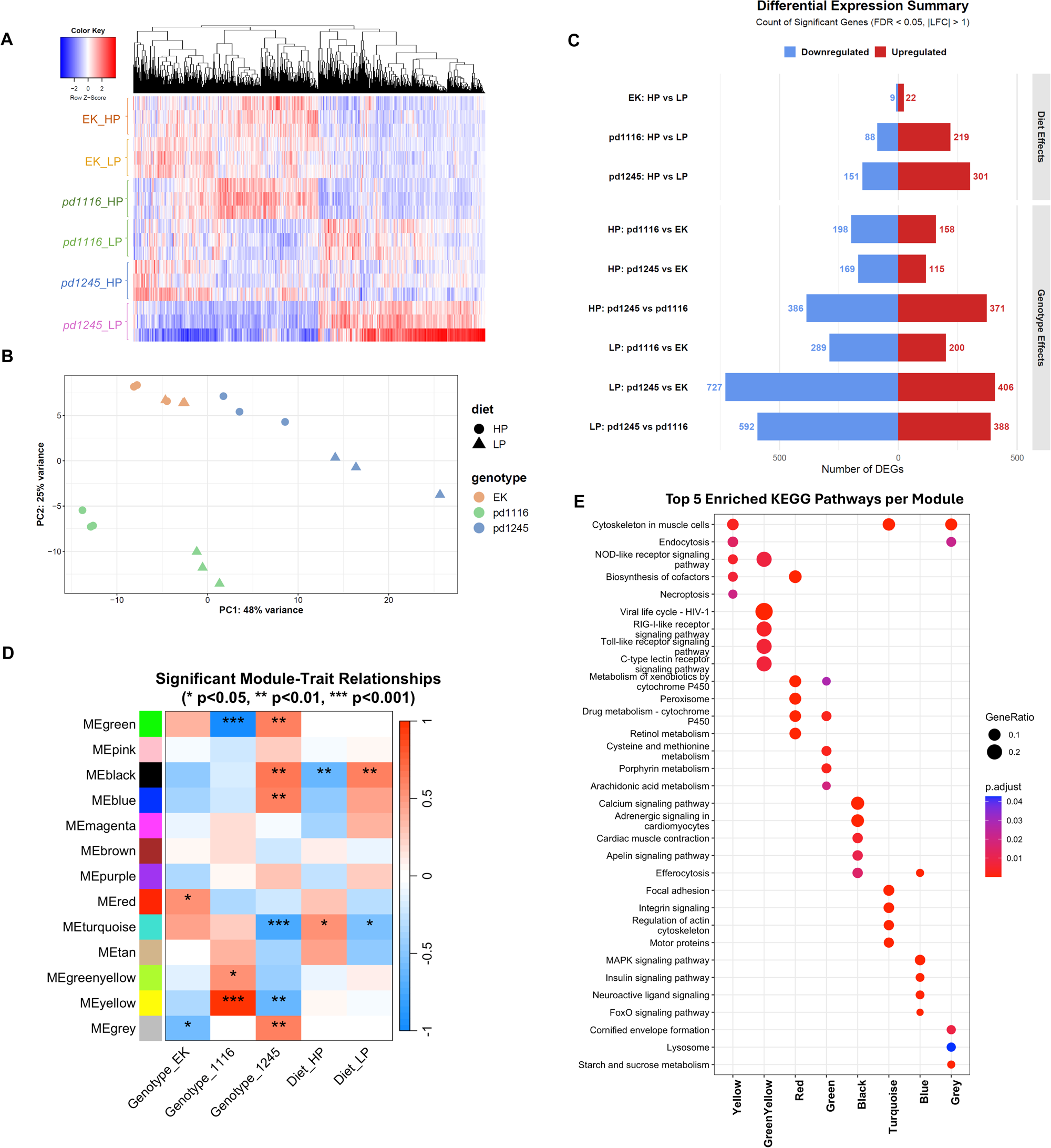
Transcriptomic analyses of WT and *pllp* alleles raised with HP and LP diets. Weighted gene co-expression network analysis (WGCNA) reveals genotype-specific transcriptional signatures. **(A)** Heatmap of all expressed genes across experimental groups, showing distinct transcriptional clustering patterns by genotype and diet. Rows represent genes; columns represent samples grouped by condition. Color scale indicates relative expression level (Z-score). **(B)** Principal component analysis (PCA) of global gene expression. PC1 separates genotypes, with *pd1116-/-* positioned closer to EK than *pd1245-/-*, reflecting transcriptional normalization. EK samples cluster together regardless of diet, while *pd1116-/-* and *pd1245-/-* show diet-dependent separation along PC1 and genotype-by-diet interaction along PC2. **(C)** Summary of differentially expressed genes (DEGs; adjusted p < 0.05, |log2FC| > 1) across all pairwise comparisons. Bar lengths indicate DEG counts; colors distinguish upregulated (red) and downregulated (blue) genes. EK shows minimal diet-responsive DEGs (31), while *pd1245-/-* under LP shows the largest transcriptional disruption (1,133 DEGs vs. EK). See also supplemental table 1. **(D)** Module-trait relationship heatmap showing Pearson correlations between module *eigengenes* and experimental traits (genotype: EK, *pd1116-/-*, *pd1245-/-*; diet: HP, LP). Color scale indicates correlation strength (red, positive; blue, negative). Four modules of interest are Yellow (positively correlated with *pd1116*, p = 3.77 × 10⁻¹²), Green (negatively correlated with *pd1116-/-*, p = 4.99 × 10⁻¹⁴), Black (positively correlated with *pd1245-/-* and LP diet), and Turquoise (negatively correlated with *pd1245-/-*, positively correlated with HP diet). **(E)** Top 5 enriched KEGG pathways for each significant module. Dot size indicates gene count; color indicates adjusted p-value; all modules included show significance in module-trait relationships.

Differential expression analysis identified markedly different numbers of differentially expressed genes (DEGs) across comparisons (Figure 2C and Supplementary Table 1). Within each genotype, EK showed only 31 DEGs between HP and LP diets (22 upregulated, 9 downregulated), confirming that diet minimally perturbs wild-type transcription. In contrast, *pd1116*-/- showed 307 diet-responsive DEGs (219 up, 88 down) and *pd1245*-/- showed 452 (301 up, 151 down), suggesting that *pllp* loss sensitizes larvae to dietary perturbation. When comparing mutants to EK, *pd1116*-/- showed 356 DEGs under HP (158 up, 198 down) and 489 under LP (200 up, 289 down), while *pd1245*-/- showed 284 DEGs under HP (115 up, 169 down) but 1,133 under LP (406 up, 727 down), which represents a dramatic expansion of transcriptional disruption under nutritional stress. The direct comparison of *pd1245*-/- versus *pd1116*-/-yielded 757 DEGs under HP and 980 under LP, underscoring the substantial transcriptional divergence between adapted and non-adapted mutants.

To identify genotype-specific transcriptional signatures, we examined the overlap of diet-responsive and genotype-responsive DEGs across conditions. Comparison of diet effects (HP vs. LP) across genotypes revealed that the majority of EK diet-responsive DEGs were conserved in the mutant backgrounds: out of the 31 EK-specific DEGs in HP vs LP, 19 (61%) overlapped with *pd1116*-/- diet-responsive genes, and 11(35%) overlapped with *pd1245*-/- (Figure S2A and Supplementary Table 2). This indicates that *pllp* mutants retain the core wild-type transcriptional response to dietary protein restriction while acquiring hundreds of additional genotype-specific diet-responsive genes expression changes (197 unique to *pd1116*-/-; 350 unique to *pd1245*-/-), reflecting a dramatically expanded sensitivity to nutritional perturbation. The larger overlap between EK and *pd1116*-/- than between EK and *pd1245*-/- is consistent with *pd1116*-/- transcriptional normalization toward the WT state. Comparison of genotype effects within each diet condition revealed distinct and overlapping gene sets between *pd1116*-/- and *pd1245*-/- relative to EK (Figure S2B, C). Under HP conditions, 237 genes were uniquely dysregulated in *pd1245*-/- versus EK but not in *pd1116*-/- versus EK, representing a potentially pathological profile specific to the non-adapted mutant. Under LP conditions, this *pd1245*-unique gene set expanded substantially, consistent with the exacerbated survival defect of *pd1245*-/- under nutritional stress. Conversely, genes uniquely dysregulated in *pd1116*-/- but not *pd1245*-/- represent candidate adaptation-associated transcriptional changes. Notably, the number of *pd1116*-unique DEGs remained relatively stable between HP (309) and LP (334) conditions, suggesting a constitutive compensatory program that operates independently of dietary context.

### Weighted gene co-expression network analysis reveals the molecular architecture of *pd1116*-/-adaptation

To capture coordinated transcriptional programs, we performed weighted gene co-expression network analysis (WGCNA), which identifies modules of co-expressed genes and correlates them with experimental traits. WGCNA identified 13 co-expression modules, six of which showed significant and biologically interpretable associations with genotype and/or diet (Figure 2D, S2D).

The Yellow module was strongly positively correlated with the *pd1116*-/- genotype, while the Green module was strongly negatively correlated. Notably, neither module showed significant association with diet (Yellow: p = 0.77; Green: p = 0.99), indicating that the transcriptional programs captured by these two modules are active in *pd1116-/-* regardless of nutritional status. This diet-independence distinguishes them from the broader transcriptional changes observed in *pd1116*-/- and suggests that these specific compensatory (Yellow) and suppressive (Green) programs have been genetically stabilized over generations of homozygous maintenance. The Red module was positively correlated with EK. Four additional modules were associated with *pd1245*-/- and diet: the Black module was upregulated in *pd1245*-/- and under LP conditions; the Blue module was upregulated in pd1245-/- without significant association to diet conditions; the Turquoise module was downregulated in *pd1245*-/- and upregulated under HP conditions, while the Grey module was upregulated in *pd1245*-/- regardless of diets (Figure 2D).

KEGG pathway enrichment of module genes revealed distinct functional signatures for each module (Figure 2E). The Yellow module (upregulated in *pd1116*-/-) was enriched for cytoskeleton in muscle cells, endocytosis, NOD-like receptor signaling, biosynthesis of cofactors, and necroptosis. Additional enriched pathways (not shown in Figure 2E) included amino acid metabolism, specifically alanine, aspartate, and glutamate metabolism, taurine and hypotaurine metabolism, and butanoate metabolism, which converge on the regulation of the glutamate-GABA axis and the GABA shunt, a metabolic bypass that utilizes glutamate decarboxylase enzymes (encoded by zebrafish *gad1a* and *gad3*) to eventually feed succinate into the TCA cycle (33). Inositol phosphate metabolism, involved in phosphoinositide signaling and vesicular trafficking regulation, was also enriched. Together, these results suggest that *pd1116*-/-adaptation involves cytoskeletal reinforcement, enhanced endocytic and vesicular trafficking, activation of immune responses and controlled cell death, and metabolic rewiring through amino acid catabolism.

The Green module (downregulated in *pd1116*-/-) was enriched for drug metabolism via cytochrome P450, porphyrin and heme metabolism, cysteine and methionine metabolism, and arachidonic acid metabolism. Suppression of cysteine and methionine metabolism, which includes the SAM-generating enzyme gene *mat2aa*, is notable because S-adenosylmethionine is the universal methyl donor for histone and DNA methylation. Additionally, enrichment for PPAR signaling, retinol metabolism, and xenobiotic metabolism indicates that *pd1116*-/- has selectively suppressed certain detoxification and lipid-sensing programs.

The Turquoise module’s eigengene expression was reduced under LP to HP diet across all three genotypes (Figure S2D). However, the magnitude of this downregulation was markedly greater in the *pllp* mutants (*pd1116* and *pd1245*). Notably, the baseline expression of this module in the *pd1245* mutants under HP conditions was suppressed to levels similar to the LP-treated *pd1116* group (Figure S2D). Functional enrichment of the Turquoise module highlighted pathways related to the muscle cell cytoskeleton, focal adhesion, integrin signaling, regulation of the actin cytoskeleton, and motor proteins (Figure 2E). This suggests that the LP condition impairs muscle structural integrity and growth, and that this physical deficit is much more severe in the *pllp* mutants than in the EK controls.

Conversely, both the Blue and Black modules exhibited increased eigengene expression under the LP diet relative to HP, a pattern that was profoundly amplified in the *pllp* mutants (Figure S2D). While the Blue module showed a dramatic upregulation in the *pd1245* LP group, the Black module demonstrated an even greater overall diet-induced shift, with the differential expression between HP and LP states vastly exacerbated in both mutant lines compared to the EK controls (Figure S2D). The Blue module was enriched for MAPK signaling, insulin signaling, neuroactive ligand-receptor interactions, and the FoxO signaling pathway, indicating a strong metabolic stress and nutrient-sensing survival response (Figure 2E) (34–37). Concurrently, the Black module was heavily enriched for the calcium signaling pathway, adrenergic signaling in cardiomyocytes, cardiac muscle contraction, and apelin signaling (Figure 2E).

Notably, both modules shared enrichment for efferocytosis. Together, these profiles imply that the LP diet provokes severe systemic stress—triggering compensatory metabolic cascades, the clearance of apoptotic cells, and massive disruptions to calcium and cardiovascular signaling—and that the loss of Pllp function leaves these animals highly vulnerable to dietary-induced physiological stress.

### *pllp* loss of function disrupts lipid metabolism, extracellular matrix integrity, and cell cycle progression

To identify the biological pathways disrupted by the loss of *pllp* function independently of adaptation, we performed KEGG pathway analysis on DEGs unique to *pd1245*-/- (non-overlapping with *pd1116*-/-) relative to EK, under both HP and LP conditions (Figure S3A). By excluding DEGs shared between both mutants which represent *pllp*-associated changes that persist regardless of adaptation, this strategy enriches for pathways that are disrupted in the non-adapted *pd1245*-/- but have been restored to near-normal expression levels in *pd1116*-/-.

Genes uniquely downregulated in *pd1245*-/- under LP conditions showed strong enrichment for vitamin digestion and absorption, steroid biosynthesis, phagosome, cell cycle, and ECM-receptor interaction pathways. Under HP conditions, uniquely downregulated genes in *pd1245*-/- were enriched for biosynthesis of unsaturated fatty acids, fatty acid metabolism, and cytoskeleton in muscle cells (Figure S3A, B). Together, these results indicate that *pllp* loss leads to a collapse of lipid and cholesterol metabolism, disruption of extracellular matrix organization, and inhibition of cell proliferation, with the severity of these defects amplified under LP conditions (Figure S3A).

To examine gene expression patterns of the lipid related DEGs (unique to *pd1245*-/- vs EK) across all experimental groups, we visualized the expression of some of the DEGs in these terms via heatmap (Figure S3B). Genes involved in lipid and cholesterol metabolism as well as vitamin digestion and absorption, including the endocytic receptor genes cubulin (*cubn*) and amnionless (*amn*), which mediate nutrient uptake in LREs (3), were markedly downregulated in *pd1245*-/- particularly under LP conditions. In *pd1116*-/-, however, these genes were largely normalized to EK levels under HP conditions, with moderate downregulation under LP that was substantially less severe than in *pd1245*-/-. A similar pattern of *pd1245*-specific downregulation with *pd1116*-/- compensation was observed for cell cycle regulators and ECM components. These results suggest that the pathways most disrupted by acute *pllp* loss, including lipid metabolism, ECM integrity, and cell proliferation, are precisely the processes that *pd1116*-/- has restored through its adaptive transcriptional program.

### *pd1116-/-* exhibits enhanced expression levels of LRE endocytic machinery genes and protein uptake kinetics

To investigate potential cell type changes linked to the adaptation of *pd1116*-/-, we first examined the expression of well-known genetic markers of various intestinal cell groups (38).Interestingly, at 20 dpf following the feeding assay, *pd1116*-/- did not show consistent upregulations of markers for intestinal cell types important for nutrient sensing or absorptions including absorptive enterocytes, LREs and enteroendocrine cells compared to EK or *pd1245*-/- (Figure S4A). We next quantified the number of active LREs by gavaging dextran 568 and segmenting the vacuoles in Ilastik and found that *pd1116*-/- did not develop more active LREs compared to EK or *pd1245*-/- (Figure S4B-D. Interestingly, *pllp* LOF in *pd1245*-/- did not seem to disrupt LRE differentiation as evidenced by the comparable vacuole number to EK (Figure S4D). Indeed, in our KEGG pathway analysis, we detected upregulation of Lysosome genes (Grey module) in *pd1245*-/- regardless of diet (Figure 2D), suggesting that reduced protein uptake upregulates lysosome biogenesis in this allele.

As mentioned above, our WGCNA and KEGG pathway analyses indicate a strong positive correlation of genes involved in endocytosis with *pd1116*-/-, including *amn* and *cubn*, which encode components of the scavenger receptor complex Cubulin-Amionless. These upregulated genes suggested a role for increased LRE function in *pd1116*-/-’s adaptation from a malnourished state. Indeed, in *pd1116*-/- many genes associated with endocytic machinery, proteases and lysosome hydrolases have comparable expressions to EK but significantly higher expression levels than *pd1245*-/-, especially under LP condition (Figure 3A, B), including the voltage-sensing phospatase gene *tpte* which is also required for protein uptake in LREs (3,39,40). To test whether this observed pattern at 20 dpf is established at an earlier developmental stage before the onset of external feeding, we conducted HCR in situ hybridization against *dab2*, *cubn*, and *amn* in 6 dpf larvae. At 6 dpf, *pd1116*-/- showed expression levels of all three LRE endocytic machinery genes that were significantly higher than those of both EK and *pd1245*-/- (Figure 3C-E’’). To investigate if such upregulation of LRE endocytosis translates to increased protein absorption, we performed a protein uptake and degradation kinetics assay in 6 dpf larvae using fast degrading mTurquoise as protein cargo (Figure 4A). We found that *pd1116*-/- was able to both take up significantly more mTurquoise protein cargo within 30 min post-gavage (PG) and degrade them much faster than EK or *pd1245-/-* over a 60 min period (Figure 4B-E). Together, these data show that transcriptional upregulations of key LRE endocytic machinery genes rather than changes in LRE numbers confer a higher protein absorption capacity to the adapted *pd1116* allele.

**Figure 3.**
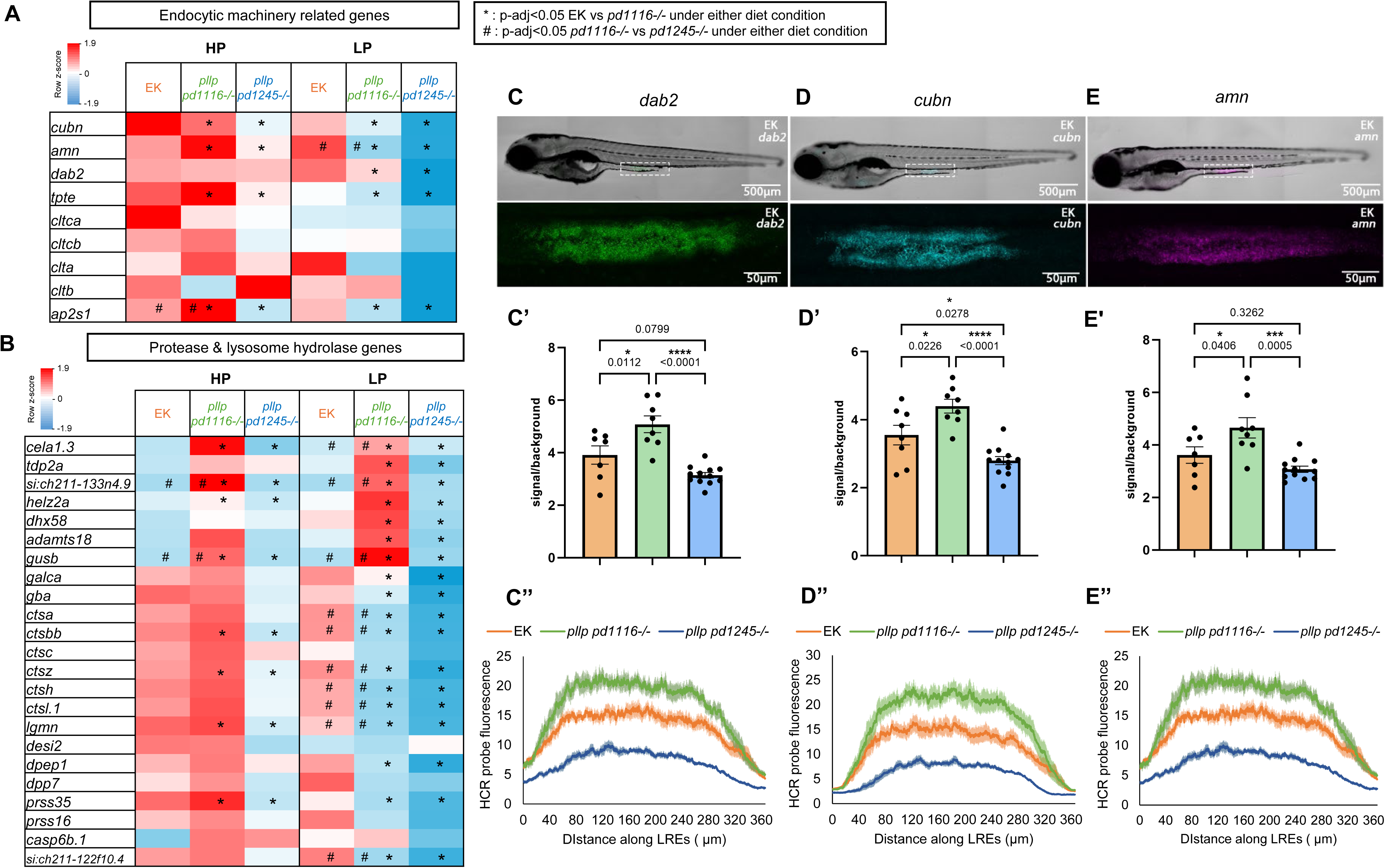
Upregulated transcription of endocytic machinery, protease, and lysosome hydrolase genes in *pd1116-/-*. **(A)** Gene expression heatmap of endocytic machinery related genes. **(B)** Gene expression heatmap of protease and lysosome hydrolase related genes. **(C-E’’)** Representative images, signal profile and expression level quantification of HCR in situ hybridization for *dab2* (C-C’’), *cubn* (D-D’’) and *amn* (E-E’’); n=7 for EK, n=8 for *pd1116-/-,* n=12 for *pd1245-/-*; one-way ANOVA.

**Figure 4.**
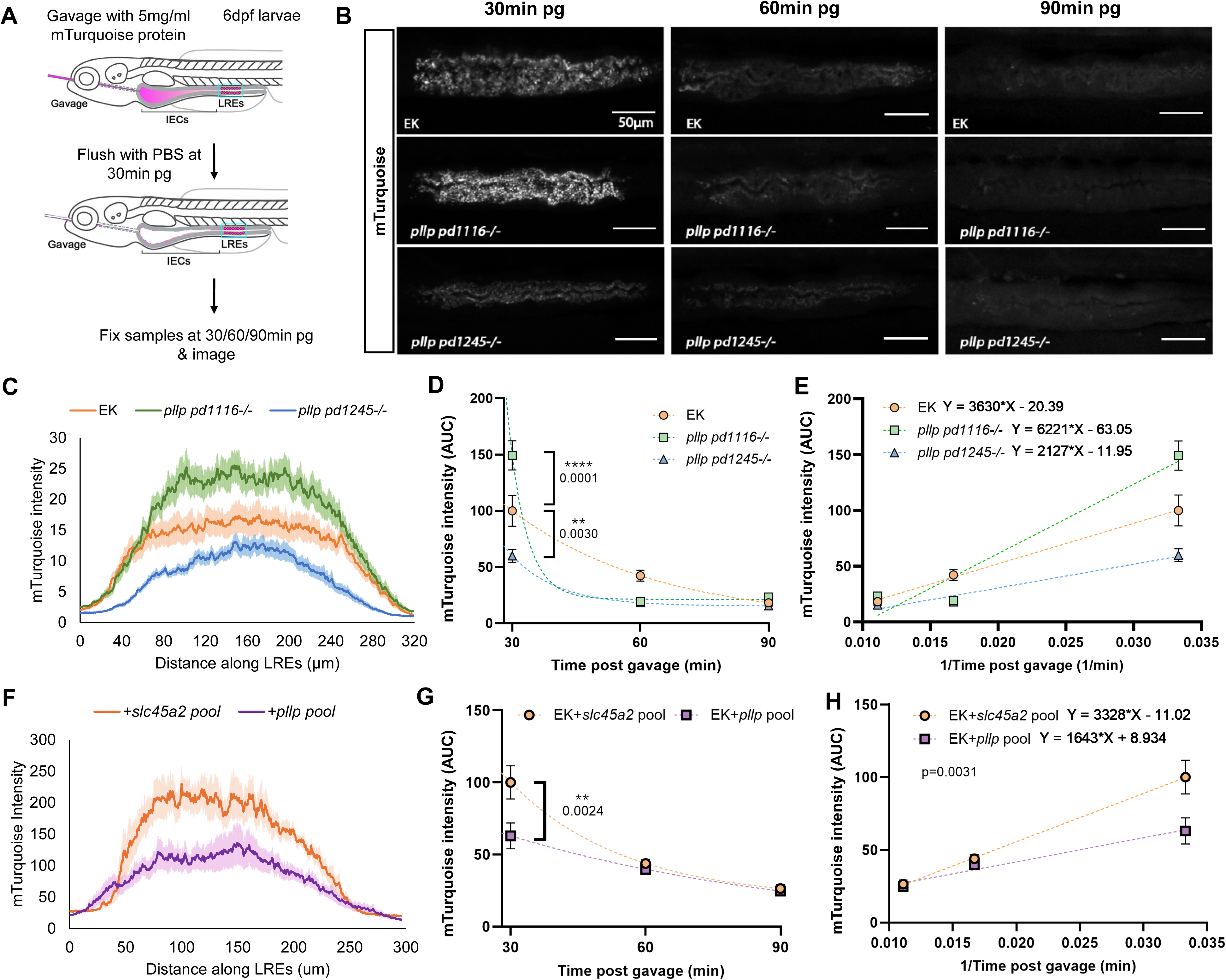
*pd1116-/-* shows enhanced protein uptake and degradation kinetics. **(A)** mTuquoise protein gavage assay experimental design; experiment were performed in 6 dpf EK, *pd1116-/-,* and *pd1245-/-* larvae. **(B)** Representative images of the LRE region at 30/60/90min post gavage (pg). **(C)** mTurquoise fluorescent signal profile at 30min pg. **(D, E)** Quantifications of mTurquoise fluorescent level curves at 30/60/90min pg (D) and linearly transformed curves showing degradation rates of each genotype (E); 30min pg: n=15 for EK, n=16 for *pd1116-/-*, n=17 for *pd1245-/-*; 60min pg: n=15 for EK, n=15 for *pd1116*-/-, n=13 for *pd1245-/-*; 90min pg: n=13 for EK, n=14 for *pd1116*-/-, n=14 for *pd1245-/-;* data normalized to EK 30min pg; 2-way ANOVA & Tuky’s test (D) ANCOVA: p<0.001 (*pd1116-/-* vs *pd1245-/-*), p<0.001 (*pd1116-/-* vs EK), p=0.2728 (*pd1245-/-* vs EK) (E). **(F)** mTurquoise fluorescent signal profile showing decreased uptake in *pllp* gRNA pool CRISPR respect to *slc45a2* control gRNA pool EK at 30min pg; n=8 for EK+*slc45a2* pool gRNA, n=10 EK+*pllp* pool gRNA. **(G, H)** Quantifications of mTurquoise fluorescent level curves at 30/60/90min pg (G) and linearly transformed curves showing degradation rates of each genotype (H); 30min pg: n=8 for EK+*slc45a2* pool gRNA, n=10 for EK+*pllp* pool gRNA; 60min pg: n=8 for EK+*slc45a2* pool gRNA, n=8 for EK+*pllp* pool gRNA; 90min pg: n=8 for EK+*slc45a2* pool gRNA, n=8 for EK+*pllp* pool gRNA; data normalized to EK+*slc45a2* pool gRNA 30min pg; 2-way ANOVA & Tuky’s test (G); ANCOVA (H).

Our quantitative protein absorption assay confirmed that loss of Pllp function in *pd1245*-/- leads to reduced protein absorption by LREs. To support this finding further, we generated a CRISPR-Cas9 gRNA pool consisting of 4 gRNAs targeting exon1-3 of *pllp,* and a control gRNA pool targeting of the pigment gene *slc45a2*(41). We validated the knock down (KD) efficiency of the *pllp* CRISPR pool by injecting into heterozygous *TgBAC(pllp^pd1114^-GFP)*, a bacterial artificial chromosome (BAC) reporter line that expresses Pllp-GFP under the control of the *pllp* regulatory sequences (Figure 1A). This resulted in a complete loss of GFP+ larvae at 6dpf as opposed to the 43.96% population of GFP+ larvae in non-injected control (NIC) group (Figure S5A). We also validated the *slc45a2* gRNA pool as a control by comparing the body length and mTurquoise uptake capability at 30min pg to NIC EK. *slc45a2* injected larvae showed an almost complete loss of pigmentation, as well as body length and mTurquoise uptake levels comparable to NIC EK, showing that *slc45a2* KD is a suitable control for LRE experiments (Figure S5B-E). Injection of Cas9 gRNA and the *pllp* gRNA pool into EK led to a significant reduction in both protein uptake and degradation kinetics compared to *slc45a2* KD controls (Figure 4F-H).

We then created a new *pllp* LOF allele, *pd130*9, using a single CRISPR gRNA targeting TM2. *pd1309* carries an in-frame deletion of 21 bp in exon 2, resulting in a 35% shorter TM2, thereby disrupting membrane insertion (Figure S5F, G). Upon assaying protein absorption, we found that the *pd1309-/-*larvae showed a significant reduction in mTurquoise protein level at 30 min PG compared to EK, similar to what we observed in *pd1245*-/- and the *pllp* CRISPR-Cas9 pool KD group (Figure S5H, I). These data demonstrate that either acute loss of *pllp* function (KD), non-sense mutant alleles (*pd1245*), or mutated alleles not subject to NMD (*pd1309*) can reproduce the protein absorption deficits of the original *pllp* mutation *pd1116* before its adaption.

Taken together, our data firmly establishes a role for *pllp* in LRE function and suggests that *pd1116*-/-developed improved LRE protein absorption capacity. This LRE hyperactivity may allow it to maximize the utilization of dietary protein under restrictive conditions, crucially contributing to its generational adaptation to protein malnutrition.

### *pd1116* adaptation involves fine-tuned expression of innate and adaptative immune response genes

Further examination of our RNA sequencing data GO enrichment on *pd1116*-specific DEGs under either diet condition revealed transcriptional changes in genes associated with response to bacterium as well as immune response (Figure 5A, B). Under HP condition, immune response DEGs had the highest relative expressions in *pd1116-/-* compared to EK, while they were downregulated in *pd1245*-/- (Figure 5A). The lowered relative expressions of DEGs in the same category in *pd1245*-/- than EK remained true under LP condition. Meanwhile, DEGs associated with response to bacterium were lower in *pd1116*-/- than EK under both diet conditions. These observations suggest that *pd1116*-/- may have become less reactive to bacterial stimuli, while still remaining immunologically active.

**Figure 5.**
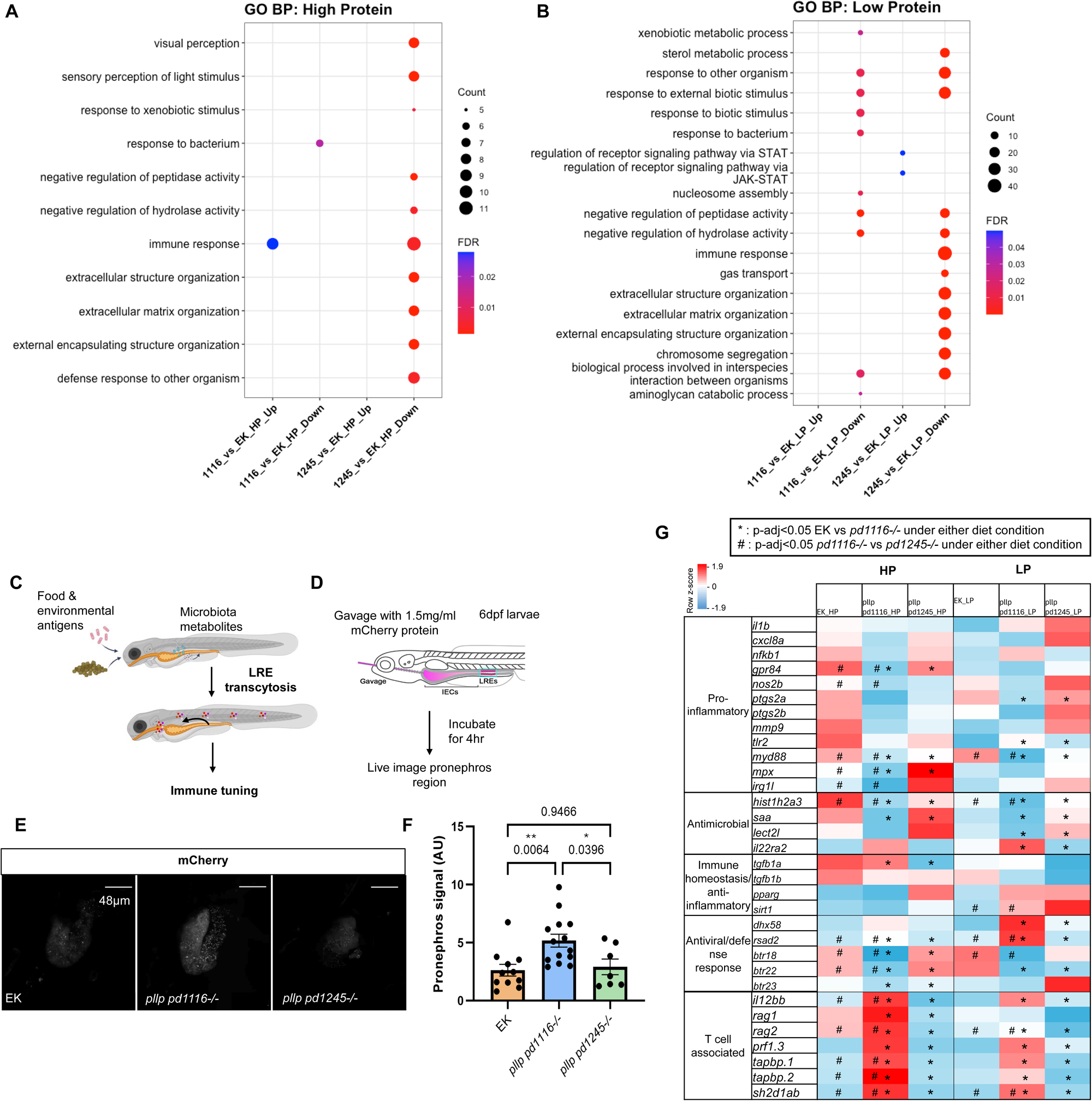
Innate and adaptative immune response adaptations in *pd1116-/-.* **(A)** GO term Biological Processes analysis results under HP feeding conditions showing DEGs upregulated (EK vs *pd1116*-HP Up) or downregulated (EK vs *pd1116*-HP Down) in EK compared to *pd1116*, and DEGs upregulated (EK vs *pd1245*-HP Up) or downregulated (EK vs *pd1245*-HP Down) in EK compared to *pd1245*. **(B)** GO term Biological Processes analysis results under LP feeding conditions showing DEGs upregulated (EK vs *pd1116*-LP Up) or downregulated (EK vs *pd1116*-LP Down) in EK compared to *pd1116*, and DEGs upregulated (EK vs *pd1245*-LPUp) or downregulated (EK vs *pd1245*-LP Down) in EK compared to *pd1245*. **(C)** Cartoon showing the proposed model of how food and environmental antigens as well as microbiota metabolites could trigger immune tuning via LRE transcytosis. **(D)** mCherry transepithelial transcytosis experiment setup. **(E, F)** Representative images of the pronephros region mCherry signal at 4h pg for EK, *pd1116-/-* and *pd1245-/-* (F) and quantifications of the signals (G); n= 11 for EK, n=14 for *pd1116-/-,* n=7 for *pd1245-/-;* one-way ANOVA. **(G)** Gene expression heatmaps showing DEGs associated with terms immune response, response to bacterium, and other well-documented genes involved in immune regulations.

Previously we documented LRE-mediated transcytosis of orally gavaged mCherry protein and transport via circulation to the pronephros, an kidney-like organ in zebrafish, via circulation (3,42). This phenomenon was particularly intriguing as it was reminiscent of the well-documented luminal antigen transcytosis process by M cells, enterocytes and goblet cells in mammals that helps balance immune activation with tolerance (43–45). Based on the heightened LRE protein uptake level and the more active immune responses in *pd1116*-/-, we hypothesize that *pd1116*-/- might have fine-tuned its immune system to better accommodate increased level of foreign antigens transcytosed by LREs (Figure 5C). To test whether *pd1116*-/- exhibits increased transcytosis, we gavaged 6 dpf larvae with mCherry protein cargo and quantified the signal intensity in the pronephros(Figure 5D). At 4 h PG, *pd1116*-/- exhibited significantly more mCherry signal in the pronephros than both EK and *pd1245*-/- (Figure 5E, F).

Close examination of DEGs within GO term pathways for immune responses and response to bacterium, as well as well-known immune genes revealed an intricate balance of immune system changes (Figure 5G). in *pd1116*-/-, we observed a general suppression of pro-inflammatory genes, downregulation of antimicrobial responses, and careful balancing of anti-inflammatory and anti-viral defense response reactions in *pd1116*-/- compared to EK or *pd1245*-/-, especially under HP condition. Interestingly, these gene regulation patterns also suggest a possible premature onset of adaptive immune maturation by 20 dpf, earlier than the literature-documented 28-42 dpf (46). This is supported by the significant upregulation of several T-cell function associated genes including *il12bb*, *rag1*, *rag2*, *prf1.3, tapbp.1, tapbp.2,* and *sh2d1ab* in *pd1116*-/- compared to EK or *pd1245*-/- under HP condition (Figure 5G).

Combined with our previous results of LRE hyperactivity, we reason that this immune response may play a role in preventing inappropriate inflammatory reactions to internalized protein macromolecules or bacterial metabolites. Such regulation may be particularly critical under malnourished conditions, as inflammations are generally considered energetically costly (47–49).

Together, these data suggest that exacerbated immune activity and inflammation may play a role in the demise of *pd1245*-/- fish, particularly under LP conditions. Conversely, a better tuned immune response in *pd1116*-/- may allow them to meet their nutritional needs while avoiding excessive inflammation caused by increased trafficking of microbial and dietary antigens. This may enable them to tolerate the significant increase in transepithelial transport across the intestine,, likely as a byproduct of increased endocytosis in LREs.

### *pllp* mutant adaptation to protein malnutrition is gradual and recurrent

To test whether we could recover the impaired LRE uptake phenotype observed in the original pre-adapted *pd1116*-/- fish, we outcrossed our current *pd1116*-/- stock to the WT AB strain (*pd1116*-AB+/-) which carries different genetic background but shows similar protein absorption capability to EK (Figure 6A,C) (50). We outcrossed *pd1116*-/- for 4 consecutive generations to AB, and in F2-F4 *pd1116*-AB+/-tested the offspring of sibling in-crosses for their protein uptake capability. Remarkably, protein absorption levels in the tested offspring populations of F2-F4 *pd1116*-AB+/-, the *pd1116*-AB-/-larvae did not revert to the pre-adapted impaired protein uptake phenotype but rather appeared to normalized with comparable uptake level to their *pd1116*-AB+/+siblings (Figure 6B). In parallel, we also performed a one generation outcross of *pd1116*-/-to AB to test if we could segregate the increased protein uptake phenotype (Figure 6A). A *pd1116*-/-parent whose offspring from a sibling in-cross showed enhanced mTurquoise absorption at 30 min PG (Figure 6C) was outcrossed to an AB parent to generate F1 *pd1116*-AB+/-. We then in-crossed the F1 *pd1116*-AB+/- adults and assayed 3 different clutches of mixed larvae and then genotyped them. Similar to previous results, we observed no significant differences in mTurquoise signal level at 30 min PG among *pd1116*-AB+/+, +/- and -/- offspring across all 4 different pairs of F1 parents we tested. These results indicate that LRE hyperactivity can be lost by outcrossing the inbred line to a different genetic background (Figure 6D). However, we did not detect a simple pattern of inheritance that could be attributed to one gene modifier.

**Figure 6:**
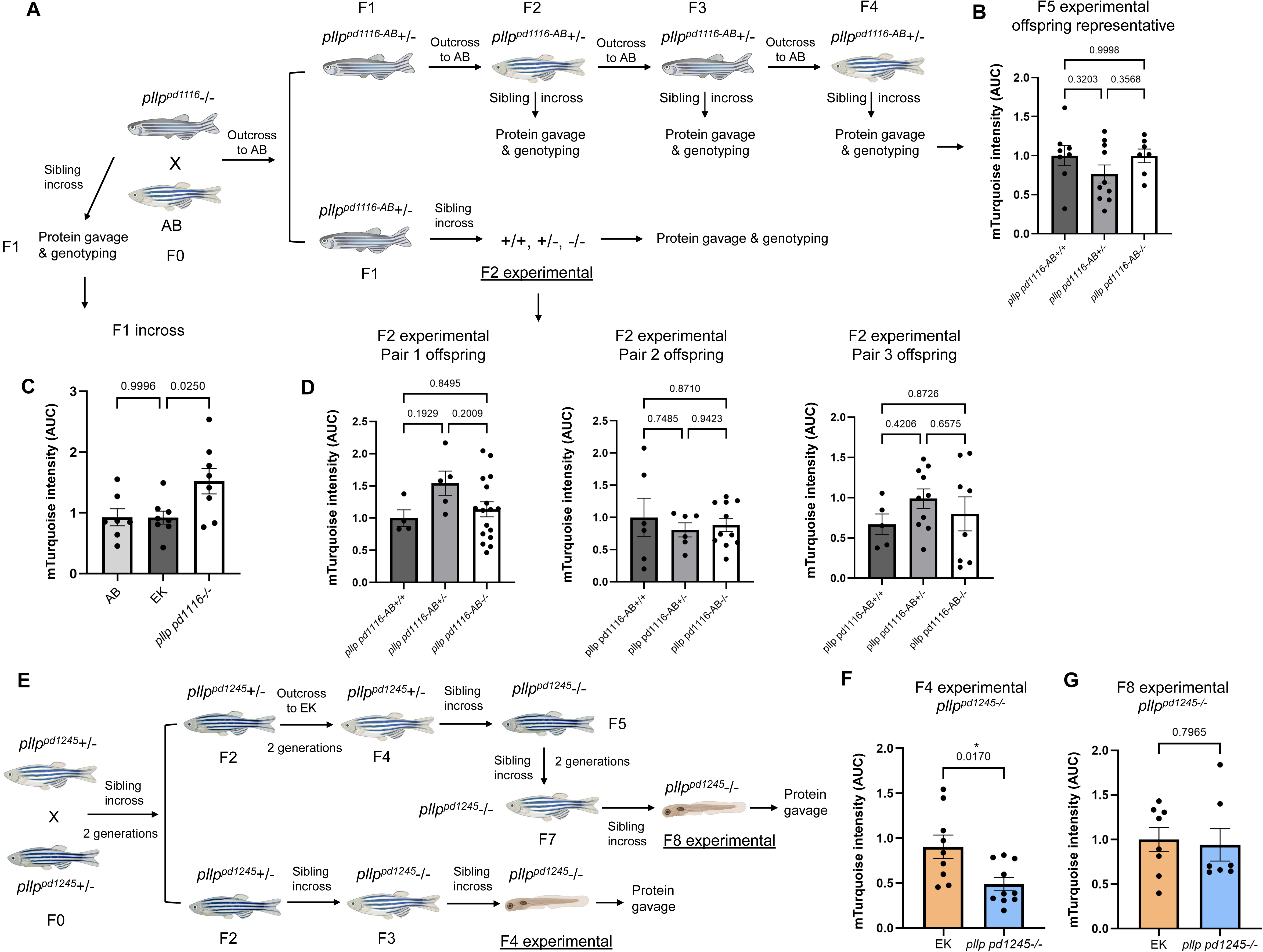
Complex and recurrent adaptation of *pllp* mutant alleles. **(A)** Pedigree of *pd1116-/-* outcrossed to unrelated WT AB; top: 3 pairs of F2 *pd1116-AB*+/- sibling in-crosses ere tested; 5 pairs of F3 *pd1116-AB*+/- sibling in-crosses were tested; 6 pairs of F4 *pd1116-AB*+/- sibling in-crosses (F5 experimental larvae) were tested. Bottom: 4 pairs of F1 *d1116-AB*+/- sibling in-crosses (F2 experimental larvae). **(B)** Representative quantification of F5 experimental offspring mTurquoise uptake level at 30min pg (from F4 *pd1116-AB+/-*bling incross) showing normalization of -/- uptake capability to +/+; n=8 for +/+, n=10 for +/-, n=7 for -/-. **(C)** 30min pg mTurquoise protein uptake level comparison between the fspring of EK, AB and the *pd1116-*/- parents used for outcrossing; n=8 for EK; n=7 for AB; n=8 for *pd1116-*/-. **(D)** Quantifications of LRE region mTurquoise protein level at 30 min pg in different clutches of F2 offspring of F1 *pd1116-AB*+/- sibling in-cross; data normalized to *pd1116-AB*+/+; one-way ANOVA; increased uptake of protein in *pd1116-/-* compared to WT is st in outcrosses but reduced uptake is not recovered even after several outcrosses. **(E)** Pedigree of the *pd1245* allele lineage; loss of reduced mTurquoise uptake level was observed arting at the in-cross experimental offspring of F5 *pd1245-/-* parents (2 in 3 clutches showed normalized protein uptake). **(F, G)** Quantifications of LRE region mTuquoise fluorescent vel at 30min pg in 6dpf F2 *pd1245-/-* experimental in-cross larvae; n=9 for EK, n=10 for *pd1245-/-* (F) and 6dpf F3 *pd1245-/-* experimental in-cross larvae; n=9 for EK, n=7 for *pd1245-.* (G); unpaired t-test; data normalized to EK.

To test whether the adaptation observed in *pd1116*-/- is recurrent, we in-crossed *pd1245*+/- according to the pedigree shown in Figure 6E. Apart from maintaining the genetic line as depicted, we also tested the protein absorption capability of the offspring of sibling in-crosses from each generation (experimental offsprings). Interestingly, we started to notice a normalization to EK levels of protein uptake levels in *pd1245*-/- starting in F6 in-cross experimental offspring (2/3 clutches), with normalization increasing in F7 incross experimental offsprings (4/5 of clutches) and becoming complete in F8 experimental *pd1245*-/-larvae (Figure 6F, G). It is worth noting that during the early maintenance of the *pd1245* allele, the *pd1245* founder heterozygous adults had been in-crossed for 2 generations before they were moved to an outcross scheme for future generations, possibly contributing to the normalization of the *pd1245* allele to EK level in our genetic experiment.

Together, these data reveal that the adaptation to the loss of *pllp* function is gradual, becoming stable, and occurring recurrently. Altogether, our studies suggest that broad transcriptional changes lead to increased protein absorption and a fined tuned regulation of immune genes that collectively result in the adaptation of *pd1116*-/- from a protein malnutrition state to one with normalized physiology that is transmitted across generations.

## Discussion

In this study, we report that zebrafish *pllp* LOF mutants overcome severe protein malnutrition due to impaired LRE function by mounting a robust and heritable transcriptional adaptation response that progressively rescues survival and growth over successive generations of homozygous selection. Through systematic comparison of the long-adapted *pd1116*-/- line with a newly generated, non-adapted *pd1245* allele and parental WT EK, we identified two intertwined adaptive mechanisms: 1-Transcriptional upregulation of the endocytic machinery in LREs, resulting in a hyperactivated protein-absorptive state that even exceeds WT capacity; 2-A coordinated, diet-independent remodeling of immune gene expression that simultaneously dampens energetically costly and potentially damaging inflammatory responses while accelerating the maturation of targeted adaptive immunity. Together, these findings reveal a previously uncharacterized mode of system-level genetic adaptation driven by persistent selection pressure, in which plasticity of both absorptive and immune function acts in concert to overcome a near-lethal genetic deficit in nutrient intake.

Remarkably, the adapted *pd1116*-/- larvae do not merely restore LRE protein absorption to wild-type levels but actually surpass it via the coordinated upregulation of several genes including *tpte*, *dab2*, *cubn*, and *amn*, thus reinforcing multiple parts of the endocytic pathway. In this context, it is notable that the similar active LRE cell numbers among *pd1116*-/-, *pd1245*-/- and EK contrast with what was suggested by earlier, largely qualitative analysis used in Rodríguez-Fraticelli et al. (2015). In this study, we quantified active LRE numbers volumetrically across the intact gut using dextran gavage and segmentation, which is a more complete and quantitative approach than the imaging of gut cross-sections used previously. Similarly, we have now assayed protein uptake quantitatively using a kinetic assay and whole-gut fluorescence signal rather than snapshots from from transverse sections, enabling more accurate comparison of total absorptive capacity across genotypes. These methodological improvements provide greater confidence that the enhanced protein absorption phenotype reported for the adapted *pd1116* allele reflects a real increase in protein uptake and degradation. Moreover, the original feeding experiment in Rodríguez-Fraticelli et al. (2015) did not adopt the calorie-restricted HP/LP feeding scheme; instead, larvae were fed a single undefined larval diet which is no longer available. This methodological difference precludes exact comparison of current survival rates to the pre-adapted state. Thus, the apparently attenuated severity of *pd1245* should be interpreted with this caveat in mind.

The second adaptive pillar involves immune regulation. Dissection of our RNAseq data and GO analyses revealed a dampening of pro-inflammatory reactions and an increased T cell function marker activation. The increased T cell activity may be due to an increase in the systemic antigen load experienced by the adapted fish due to increased transcytosis. The simultaneous suppression of pro-inflammatory gene expression in *pd1116*-/- can therefore be understood as an adaptive countermeasure. Inflammatory innate immune responses can cause tissue damage to the gut (51) and other tissues, thus tuning down inflammatory innate responses while speeding up the maturation of the more targeted adaptive immune responses may be particularly beneficial in improving survival, especially under nutrient restriction (30–32, 38,39). This is illustrated by the pronounced divergence of responses under HP conditions, precisely the dietary context in which LRE hyperactivity would increase transcytosis of luminal protein antigens and bacterial metabolites into other organs via circulation. This also suggests that the predicted decrease in sensitivity to bacteria in *pd1116*-/- from our GO term analysis may represent a protective response.

Furthermore, because the endocytic machinery is negatively regulated by microbial signals (31), reduced sensitivity to bacteria in the adapted *pd1116*-/- fish may account for the increased activity compared to WT. However, whether protein uptake in the adapted *pd1116* allele is insensitive to microbial signals is unclear.

### Metabolic rewiring in the adapted state

WGCNA revealed that *pd1116*-/- adaptation is accompanied by metabolic rewiring. The Yellow module, which is constitutively upregulated in the adapted *pd1116*-/-, is enriched not only for endocytic and immune pathways, but also for amino acid catabolism. These pathways eventually feed into the TCA cycle, potentially boosting mitochondrial energy production independently of canonical glucose and amino acid oxidation. This would place additional strain on mitochondrial function, making mitophagy dependent protective pathways crucial (54). In this context, downregulation of autophagy in *pd1245*-/-may partially explain the loss of LRE activity.

On the other hand, suppression of SAM biosynthesis genes in the adapted line raises the intriguing possibility that altered epigenetic writing capacity may itself contribute to the stabilization of the adaptive transcriptional state, by reducing the availability of methyl groups needed to silence compensatory gene programs established in earlier generations(55). Whether this represents a cause or a consequence of the broader adaptation remains to be determined, but it opens a potentially important link between metabolic state and epigenetic heritability in the context of protein malnutrition.

### Genetic origin of nutritional adaptation

While our analyses have uncovered key functional mediators of the adaptation of *pd1116*-/-, the genetic origin remains unclear. Both *pd1116*-/- and *pd1245*-/- transcripts showed nonsense-mediated mRNA decay (NMD). NMD of a mutant transcript has been shown to lead to transcriptional adaptation (TA), the compensatory upregulation of sequence-related genes, typically paralogs, via transcriptional effects driven by mRNA decay products(56–60). This response may also be epigenetically inherited across generations through germline-carried signals, with H3K4me3 enrichment at adapting gene promoters detectable in early embryos of subsequent generations (61,62). This mechanism is therefore a plausible contributor to the stable, multigenerational transcriptional changes we observe in *pd1116*-/-. However, known *pllp* paralogs including *cmtm3*, *cmtm6*, *cmtm7*, *cmtm8a*, *mal2*, *marveld1*, *plp2b*, and *malb*, did not show significant upregulation in either *pd1116*-/- or *pd1245*-/- compared to EK (Figure S6), as canonical NMD-triggered TA would predict. *cmtm3* and *cmtm8a* were upregulated in *pd1116*-/- relative to *pd1245*-/-under HP conditions, and *cmtm3* and *cmtm7* showed the same pattern under LP conditions. This differential expression between the two alleles despite both triggering NMD suggests either that *pd1116*-/-has developed a non-canonical or partial form of TA specific to certain paralog members, or that the compensation occurs through network-level or non-paralog gene upregulation. Another important difference between the adaptation observed in *pd1116*-/- and canonical TA is that while the former seems to arise gradually, the latter occurs acutely withing hours of expressing a mutated mRNA subject to NMD(56). Thus, whether NMD dependent TA contributes to the adaptation observed in *pllp* alleles is unclear. Also unclear is how a broad response involving genes of unrelated sequence and spread across the genome may be coordinated and inherited across generations.

Finally, it is worth noting that the *pd1116* allele was generated with a TALEN, while *pd1245* was generated with CRISPR/Cas9. It is possible that these two genome engineering methods alter chromatin organization and gene expression in different ways. Beyond the differences in editing modality, it is also important to recognize that the original *pd1116* mutation could not be precisely recreated due to the technical limitations of the early CRISPR-based approaches used for generating the *pd1245* allele.

However, the phenotypes of both alleles are consistent with complete loss of function, as both produce large truncations and also show pronounced *pllp* transcript degradation by RT-PCR, consistent with NMD. Nevertheless, it is possible that differences in the genomic lesion between the two alleles, or subtle differences in the residual mutant RNA fragments produced, contribute to the divergent degrees of phenotypic severity relative to what was originally reported for *pllp* loss of function.

## Materials and Methods

### Fish

The Duke University Institutional Animal Care and Use Committee (IACUC) guidelines were followed in the care and use of all fish in this project. We maintained zebrafish (*Danio rerio*) stocks on a recirculating system at 28°C with a controlled, 14-hour light and 10 hour dark cycle (). Genotypes were determined by fin clipping. Male and female breeders from 3-9 months of age were used to generate fish for all experiments. Breeding adult zebrafish were fed a 1:1 ratio of GEMMA Micro (Skretting) and artemia. To breed fish, males and females were placed in mating tanks with dividers overnight, and dividers were removed the following morning. 5-30 dpf zebrafish larvae from the Ekkwill (EK) or AB background were used in this study. No test on the influence of sex was performed as sex determination in zebrafish occurs in late juvenile stage, well after our experimental window. Strains generated for this study: *pllp^pd1116^, pllp^pd1245^, pllp^pd1309^*.

### Genome Editing

Mutant lines and CRISPR-Cas9 pool knockout fish were generated using CRISPR/Cas9. Guide RNA (gRNA) target sites were identified using CRISPRscan (63) and gRNAs were synthesized using the oligo-based method(64). *pllp pd1245* mutants were generated using a gRNA targeting exon2, and *pllp pd1309* mutants also at exon2. gRNA target sequences were: *pd1245*- 5’ GGTTGCTGAGTCTTCATCAG 3’, and pd1309- 5’ GGTTCTGAGCATTGCCCTGC 3’. gRNA target sequences for *pllp* CRISPR-Cas9 pool are: 5’ GGCTGAGTGCTGACCTTCCC 3’, 5’ GGTTCTGAGCATTGCCCTGC 3’, 5’ GGACCCTGGTAATACCCACC 3’, 5’ GGTGAAGGGCAGAACGCAGC 3’. gRNA target sequences for *slc45a2* CRISPR-Cas9 pool are: 5’ GGACTACTGTAGGTCGTCAT 3’, 5’ GGGTCCGTCAATGAAGTCTG 3’, 5’ GGATGCTAACCATAGCTGAC 3’, 5’ GGCTATGCCACCTATGAGAG 3’. Zebrafish embryos were injected at the one cell stage with 150 pg/nl of Cas9 mRNA and 50 pg/nl of gRNA.

Genotyping for *pd1116* was performed using primers: forward, 5’ GCACTCAGACCAGCTCACAA 3’; (DCAPS primer (65)) reverse, 5’ CGGAACAGAAAAGTGGGTGT 3’. PCR products were then digested with BsmI enzyme for genotyping. Genotyping for *pd1245* was performed using primers: forward, 5’ GCTGGTTATCCTGTTGCTGAT 3’; reverse, 5’ GCATTAGCCAGAAAGGCAGT 3’. Genotyping for pd1309 was performed using primers: forward, 5’ GATTATTAGTGTGGACGCTGATTGC 3’; reverse, 5’ AATGTTCCCTCCTTCATGAATAGTC 3’.

### RNA Isolation and Reverse Transcription PCR (RT-PCR)

RNA was extracted using the RNeasy Mini Kit (Qiagen) according to the manufacturer’s protocol. cDNA was synthesized using first Strand cDNA Synthesis Kit (Roche). To generate cDNA for RT-PCR, poly dT primer was used for reverse transcription cDNA synthesis. Primers used for RT-PCR are: pllp_F 5’ GGCGGATTTTCCTGGGAAGG 3’, pllp_R 5’ CCGTGCTCCCTGCTGC 3’; rpl13a_F 5’ TCTGGAGGACTGTAAGAGGTATGC 3’, rpl13a_R 5’ AGACGCACAATCTTGAGAGCAG 3’. *rpl13a* was used as standard.

### Fluorescent Protein Purification

Previously described methods were used to prepare mCherry and mTurquoise (3).

### Microscopy

Whole-mount confocal live or fixed sample imaging was performed on a 1) Leica SP8 confocal microscope with a 25X objective (Leica) and (Leica), or 2) Olympus confocal microscope with a 30x/1.05 silicone oil objective (Olympus) and FluoView software (Olympus). Fish were mounted onto glass-bottom dishes in a 0.9% agarose mixture of egg water and 1x tricaine. Z-step size for z-stacks images was 1.5 μm.

### Gavage Assays

Larvae were sedated with 0.22 µm-filtered 1X Tricaine (0.2 mg/mL). Sedated larvae were suspended in 3% methyl cellulose and gavaged with ∼4 nL of fluorescent cargo(66). Larvae were placed in a 28°C zebrafish incubator to absorb the fluorescent cargo following gavage.

### mTurquoise Uptake & Degradation

Larvae were gavaged with mTurquoise (2.5 mg/mL) and incubated for 30min. Luminal mTurquoise was cleared with a 1X PBS gavage. Following clearance, larvae were preserved in 4% PFA in PBS immediately following clearance, 30min post clearance or 60min post clearance. Samples were stored at 4° C overnight. The following day, larvae were mounted on glass bottomed dishes and imaged by confocal microscopy. Fluorescence profiles of mTurquoise signal in LREs were generated by analyzing confocal images in ImageJ (version 1.53t). Images were z-projected as max intensity plots. The LRE region was delineated with a rectangular selection tool to generate plot profiles of mTurquoise fluorescence along the LRE region. Signal intensity was calculated from area under the curve (AUC) measurements in PRISM Graphpad. The average AUC measure at each time point was calculated and plotted. These values were normalized by dividing them by the average AUC value at 30 minutes post gavage in EK.

The rate of mTurquoise degradation was calculated by doing a simple linear regression on the plotted average AUC values at 30/60/90min pg. The differences in the degradation slopes among EK and pllp mutants or CRISPR knock-out groups were calculated and compared for differences in mTurquoise degradation rates.

### mCherry Trans-Epithelial Transport

6dpf EK*, pllp ^pd1116^-/-,* and *pllp ^pd1245^-/-* larvae were sedated & gavaged with mCherry protein (1.25 mg/mL) and incubated for 4 hours. After that, sedated larvae were mounted on glass-bottom plates and live imaged by confocal microscopy. The average mCherry intensity in the pronephros organ was quantified and compared using 1-way ANOVA with Tukey’s multiple comparisons test.

### Dextran568 Gavage for LRE Segmentation

6dpf EK*, pllp ^pd1116^-/-,* and *pllp ^pd1245^-/-* larvae were sedated & gavaged with a dextran568-phenol red gavage mixture (1.5mg/mL Dextran568 in 1X PBS with 0.03% phenol red) using the same method described previously. After 1 hour of incubation, the larval gut was cleared as described above.

### LRE Segmentation

Following dextran568 gavage, larvae were stored in the zebrafish incubator for 26 hours. Larvae were then sedated and live imaged by confocal microscopy. Images were imported into Ilastik version 1.3.3 and segmented by the Pixel Classification program. The program is trained to identify dextran568-filled LRE vacuoles by machine learning where 10 – 15 LRE vacuoles are manually marked per image. This process is carried out for 5 – 10 images before the program is allowed to train itself based on these manual feed-ins. Any errors made by the program are then manually corrected and used to improve the segmentation process. After the segmentation program is trained to have at least an estimated 85% accuracy, the results are exported as probability maps in .h5 format, which are then fed into Ilastik’s Object Classification program. The following parameters are used for the classification process: Method: Hysteresis; Smooth: σ=1.0 for x-,y- and z-axis; Threshold: core=0.60, final=0.60; Don’t merge objects. After checking the accuracy of object classification, the results are then exported as .cvs files from which the numbers of objects (in this case LRE vacuoles) are extracted. Statistics were done by 1-way ANOVA with Tukey’s multiple comparisons test.

### *In Situ* Hybridization Chain Reaction (HCR)

HCR probes, hairpins, amplification, wash and hybridization buffers were purchased from Molecular Instruments(67). Our methods were adapted from a previously published procedure for performing HCR on zebrafish embryos(68). At 6dpf, larvae were anesthetized and fixed in 4% PFA, then incubated on a shaker table for two hours. Larvae were washed with 1x PBS twice, followed by two cold acetone washes. Larvae were incubated in cold acetone at −20°C for 8 minutes, followed by three 1x PBS washes. For the detection stage, larvae were first incubated in probe hybridization buffer at 37°C for 30 minutes on a shaker. Larvae were incubated in probe solution (4 nM) on a shaker at 37°C for 24 – 48 hours.

Excess probes were removed by washing larvae four times with preheated, 37°C probe wash buffer, incubating samples on a shaker table at room temperature for 15 minutes each time. This step was followed by two, 5-10 min SSCT washes on a shaker at room temperature. For the amplification stage, larvae were incubated in room temperature amplification buffer for 30 minutes. Hairpins (30 pmol) were prepared by heating at 95°C for 90 seconds, then snap-cooled in the dark at room temperature 30 minutes. Hairpins were added to amplification buffer. Larvae were incubated in the hairpin solution in the dark at room temperature overnight. Excess hairpins were removed with five SSCT washes on a shaker table at room temperature. The duration of the first two washes were 5 minutes, followed by two 30-minute washes and one 1-minute wash. Samples were protected from light during the washes. Larvae were imaged by confocal microscopy using the Olympus system. Fluorescence profiles of each samples were quantified by taking the ratio of the average signal of the LRE region divided by the average signal of the background area. The differences in quantified fluorescence profiles among larvae being compared were calculated with 1-way ANOVA with Tukey’s multiple comparisons test.

### Zebrafish Growth and Survival Under Controlled High- and Low-Protein Diet Feeding Conditions

Sibling larvae from EK, *pllp^pd1116^*-/-, and *pllp^pd1245^*-/- incrosses were conventionally raised from 0-5 dpf. At 6 dpf, 10 larvae/3L tank were placed on our recirculating Aquatic Habitats system. From 6-20 dpf, larvae were fed 10 mg/day of a custom formulated high (HP) or low-protein (LP) diet daily between 11 am-12 pm. Diet formulations are described in detail below. The survival of fish was monitored up to 20 dpf.

Then, surviving fish were collected and imaged at 20 dpf with an Axio Zoom.V16 Stereoscope (Zeiss) with ZEN pro 2012 software (Zeiss). Using the acquired images, fish standard lengths (69) were measured with ImageJ. A separate image of 10mm ruler under the same zoom parameter was taken as the size reference.

### Diet Formulations

Previously described diet formulations were used in the above feeding experiments (31).

### RNA extraction and bulk RNA sequencing and analysis

Zebrafish embryos/larvae were collected, rinsed in ultra-pure water, and immediately snap-frozen in liquid nitrogen. Tissues were homogenized into a fine powder using a pre-chilled, RNase-free disposable pellet pestle. Total RNA was extracted by adding 300 µL of TRIzol reagent and allowing the homogenate to thaw. Following the addition of 60 µL of chloroform and vigorous vortexing, the phases were separated using Phase Lock Gel Heavy tubes (QuantaBio) centrifuged at 12,000 rcf for 15 minutes at 4°C.

The aqueous phase was recovered and further purified using the RNeasy MinElute Cleanup protocol (Qiagen). Briefly, the sample was mixed with 70% ethanol and transferred to an RNeasy MinElute spin column. The membrane was washed with Buffer RPE and 80% ethanol, followed by a 5-minute centrifugal drying step with open lids to eliminate residual ethanol. RNA was eluted in 14 µL of RNase-free water. RNA concentration was evaluated by Qubit (Thermo Fisher Scientific) and integrity was assessed using an Agilent 2100 Bioanalyzer. Samples with an RNA integrity number (RIN) >7.0 were used for sequencing. Libraries were prepared using a stranded mRNA-Seq protocol and sequenced on the Illumina NovaSeq 6000 platform (S-Prime flow cell) with 50 bp paired-end reads.

Raw sequence quality was assessed using FastQC (v0.11.7) (70), and adapter sequences and low-quality bases were removed using Trim Galore (v0.6.5) (71). Trimmed reads were aligned to the Zebrafish genome (GRCz11/danRer11) and the corresponding Ensembl transcriptome (72) (Release 100) using STAR (v2.7.3a) (73). Genome indices were generated using a --sjdbOverhang of 49. For alignment, --outFilterMultimapNmax was set to 1 to retain only uniquely mapped reads, and --outSAMstrandField intronMotif was utilized for downstream compatibility.

Gene quantification was performed using HTSeq (v0.11.1) (74). Differential expression analysis was conducted in R using DESeq2 (v1.26.0) (75). Prior to normalization, genes with fewer than 10 reads in at least one condition were filtered out. The DESeq() function was used to normalize data and estimate dispersions. Differential expression was modeled using the Wald test. The results were generated without independent filtering or Cooks distance-based outlier filtering to ensure a comprehensive results table.

Genes were considered differentially expressed if they exhibited a log2 fold change ≥ 2 and an adjusted p-value < 0.05 (Benjamini-Hochberg correction (76)). For visualization, Variance Stabilizing Transformation (VST) was applied to the count data. Principal Component Analysis (PCA), MA-plots, and hierarchical clustering heatmaps (Spearman correlation distance, complete linkage) were generated using the DESeq2 and gplots (77) packages.

Functional enrichment analysis was performed on the DEGs identified from all experimental comparisons, including dietary effects within each genotype (HP vs. LP for EK, 1116, and 1245) and genotypic differences under specific diets (1116 vs. EK and 1245 vs. EK for both HP and LP conditions). Over-representation analysis for Gene Ontology (GO) Biological Process (BP) and KEGG pathways was conducted using the clusterProfiler package. To maintain statistical rigor, custom backgrounds were utilized for all tests, consisting of all genes detected in the RNA-seq dataset (22,863 genes for GO using gene symbols and 20,439 genes for KEGG using Entrez IDs). P-values were adjusted using the Benjamini-Hochberg method, with a significance threshold of adjusted p < 0.05.

### Weighted Gene Co-expression Network Analysis (WGCNA)

Weighted gene co-expression network analysis was performed using the WGCNA package (74) in R to identify modules of highly co-expressed genes. To focus the analysis on biologically informative variation, count data were first normalized using a Variance Stabilizing Transformation (VST) from the DESeq2 package (75). The top 25% most highly variable genes (based on variance) were retained for network construction, resulting in a dataset of 5,716 genes.

A signed hybrid co-expression network was constructed using a soft-thresholding power (β) of 18. Modules were identified using the blockwiseModules function with a maximum block size of 20,000 genes, a minimum module size of 30, and a merge cut height of 0.25. For each module, a Module Eigengene (ME)—representing the first principal component of the expression profile—was calculated. Module-trait relationships were evaluated by correlating MEs with experimental traits using Pearson correlation. To visualize expression patterns across genotypes and diets, module eigengene values were plotted as boxplots.Functional annotation was performed using the clusterProfiler (79) package and the org.Dr.eg.db zebrafish database. Gene Ontology (GO) enrichment analysis for biological processes (BP) and KEGG pathway enrichment were conducted for genes within specific modules. Enrichment analysis was performed using the standard organism-wide background to ensure broad capture of functional categories. P-values were adjusted for multiple testing using the Benjamini-Hochberg (BH) method, with a significance threshold of q < 0.05.Bibliography for RNAseq analysis.

## Ethics Statement

Animal husbandry and experimental procedures followed the guidelines set forth by the Association for Assessment and Accreditation of Laboratory Animal Care (AAALAC). All zebrafish experiments were performed in accordance with protocols approved by the Duke University Institutional Animal Care and Use Committee (IACUC) under protocol registry number 57 approved on February 21st, 2026.

## Acknowledgements

We thank John Rawls and members of the Bagnat lab for insightful discussions; Dan Levic and Carina Block for critical reading of our manuscript; Colin Lickwar for helpful advice on bioinformatic analysis; Akankshi Munjal and Yusuke Mori for advice on HCR; and the Duke ZeCore staff for fish care.

This work was funded by NIH grant DK303003784 to MB.

---

**Figure S1:**
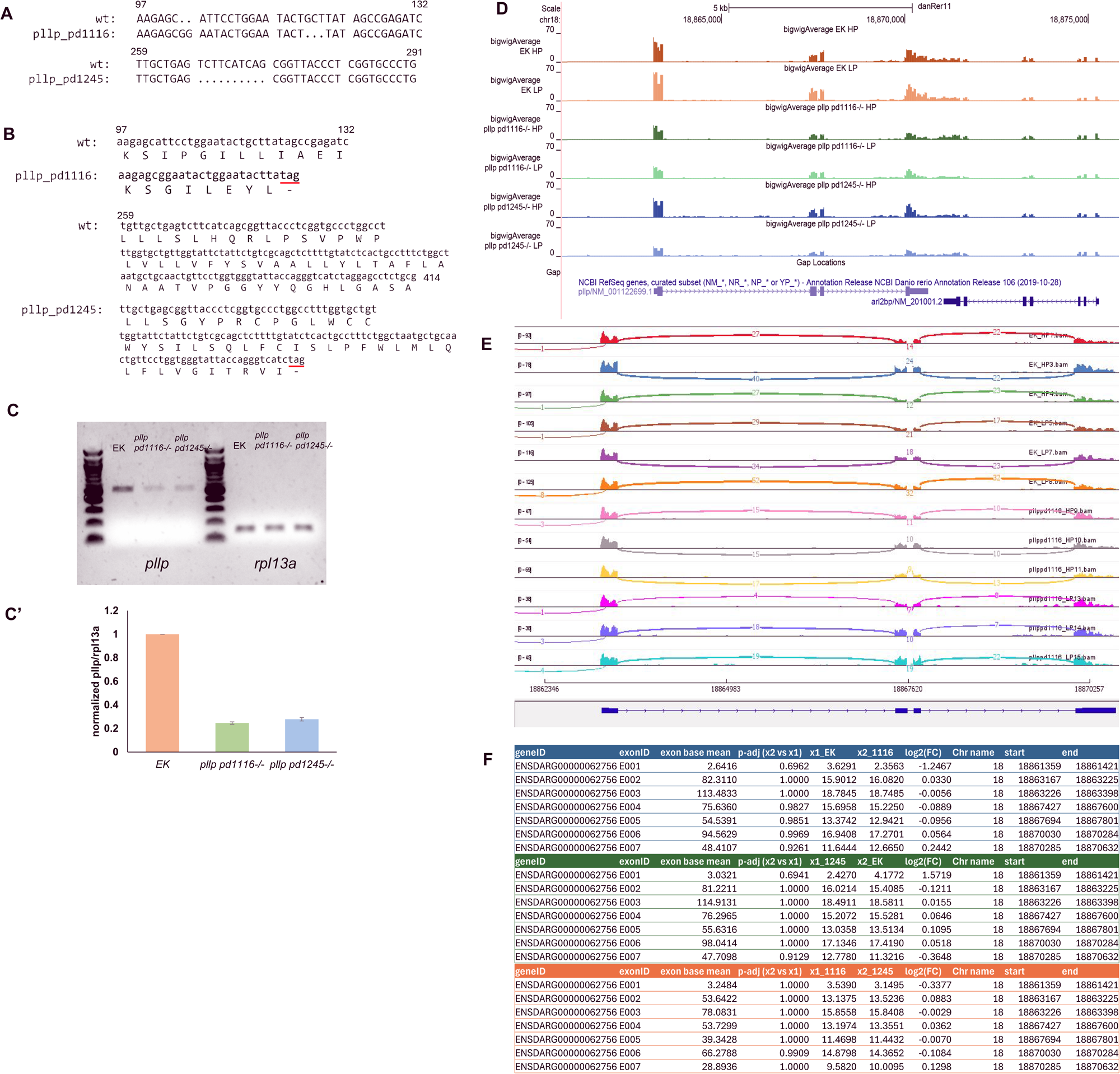
Analyses of *pllp* locus sequence and mRNA splicing. **(A)** DNA sequencing results at the original *pd1116-/-* mutation site (top) and the *pd1245-/-*mutation site (bottom); numbers indicate bp downstream of the ATG start codon. **(B)** Predicted protein sequences of Pllp^pd1116-/-^ (top) and Pllp^pd1245-/-^ (bottom); numbers indicate aa downstream of the Met start site. **(C)** bigwig file of the *pllp* locus showing tracks for EK, *pd1116-/-* and *pd1245-/-* under both diet conditions. **(D)** IGV visualization of splice junction coverage at the *pllp* locus for EK and *ppd1116-/-* under both diet conditions; no novel splicing patterns are detected. **(E)** DEX Seq analysis of relative exon usage at the *pllp* locus. **(F, F’)** RT-PCR of *pllp* and *rpl13a* control **(F)** and quantifications of band intensity **(F’)** comparing EK, *pd1116-/-* and *pd1245-/-.* Both mutant transcripts undergo non-sense mediated decay.

**Figure S2.**
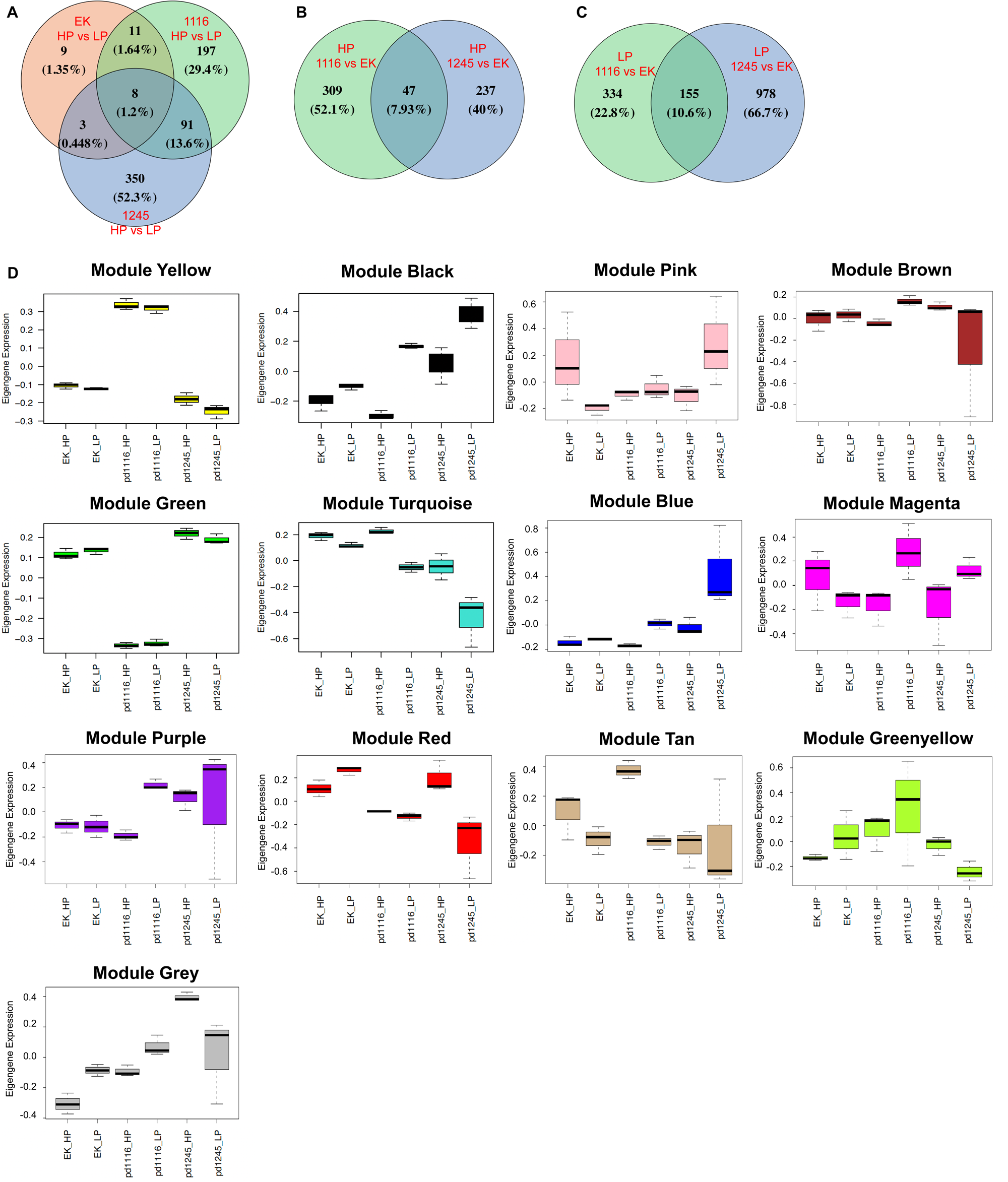
Transcriptomic profiling reveals genotype-dependent and diet-dependent gene expression programs in *pllp* mutants. **(A)** Venn diagram showing overlap of diet-responsive DEGs (HP vs. LP) among EK, *pd1116-/-*, and *pd1245-/-*, illustrating genotype-specific transcriptional sensitivity to diet. **(B)** Venn diagram of DEGs identified in HP *pd1116-/-* vs. EK and HP *pd1245-/-* vs. EK, highlighting genes uniquely dysregulated in each mutant under HP conditions. **(C)** Venn diagram of DEGs identified in LP *pd1116-/-* vs. EK and LP *pd1245-/-* vs. EK, revealing the expanded set of *pd1245*-unique DEGs under nutritional stress. See also supplemental table 2. **(D)** Eigengene expression bar plots for the all modules across all experimental groups (EK-HP, EK-LP, *pd1116*-HP, *pd1116*-LP, *pd1245*-HP, *pd1245*-LP), illustrating the genotype and diet specificity of each module. Yellow and Green modules show strong *pd1116*-specific patterns independent of diet, while Black and Turquoise modules show combined genotype and diet effects.

**Figure S3.**
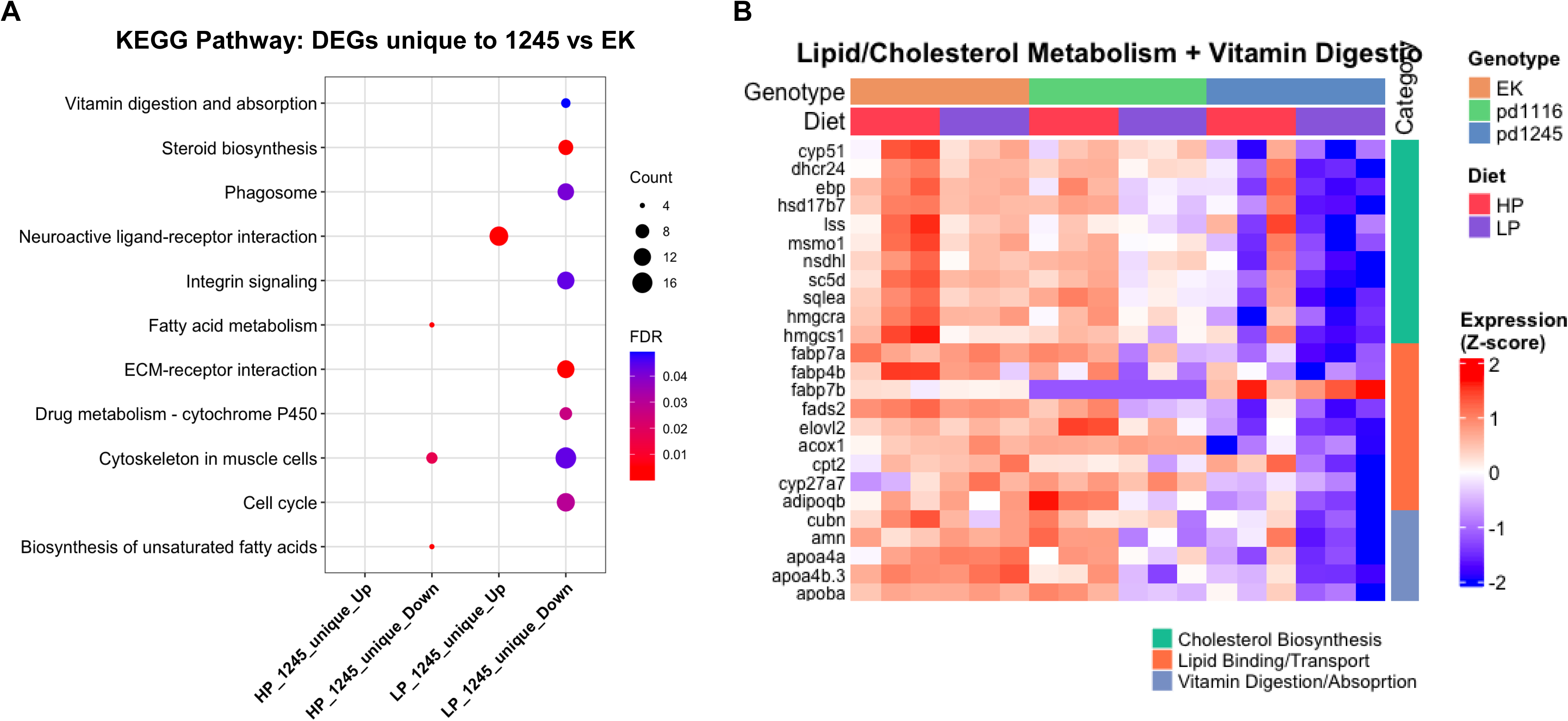
KEGG pathway analysis shows *pd1245* mutation disrupts lipid metabolism, extracellular matrix integrity, and cell cycle progression. **(A)** Comparative KEGG pathway enrichment (compareCluster) of DEGs uniquely dysregulated in *pd1245-/-* relative to EK (non-overlapping with *pd1116-/-* vs. EK DEGs). Four gene lists were analyzed (left to right): upregulated and downregulated DEGs unique to *pd1245-/-* under HP and LP conditions, respectively. Dot size indicates gene count; color indicates adjusted p-value. **(B)** Heatmap of gene expression across all six experimental groups for selected functional categories: lipid/cholesterol metabolism and vitamin digestion and absorption (including the components of LRE endocytic machinery *cubn* and *amn*). Color scale represents Z-score. In *pd1245-/-*, these genes are broadly downregulated, with the most severe suppression under LP conditions. In *pd1116-/-*, expression is largely normalized, particularly under HP conditions, demonstrating transcriptional compensation of these critical pathways.

**Figure S4.**
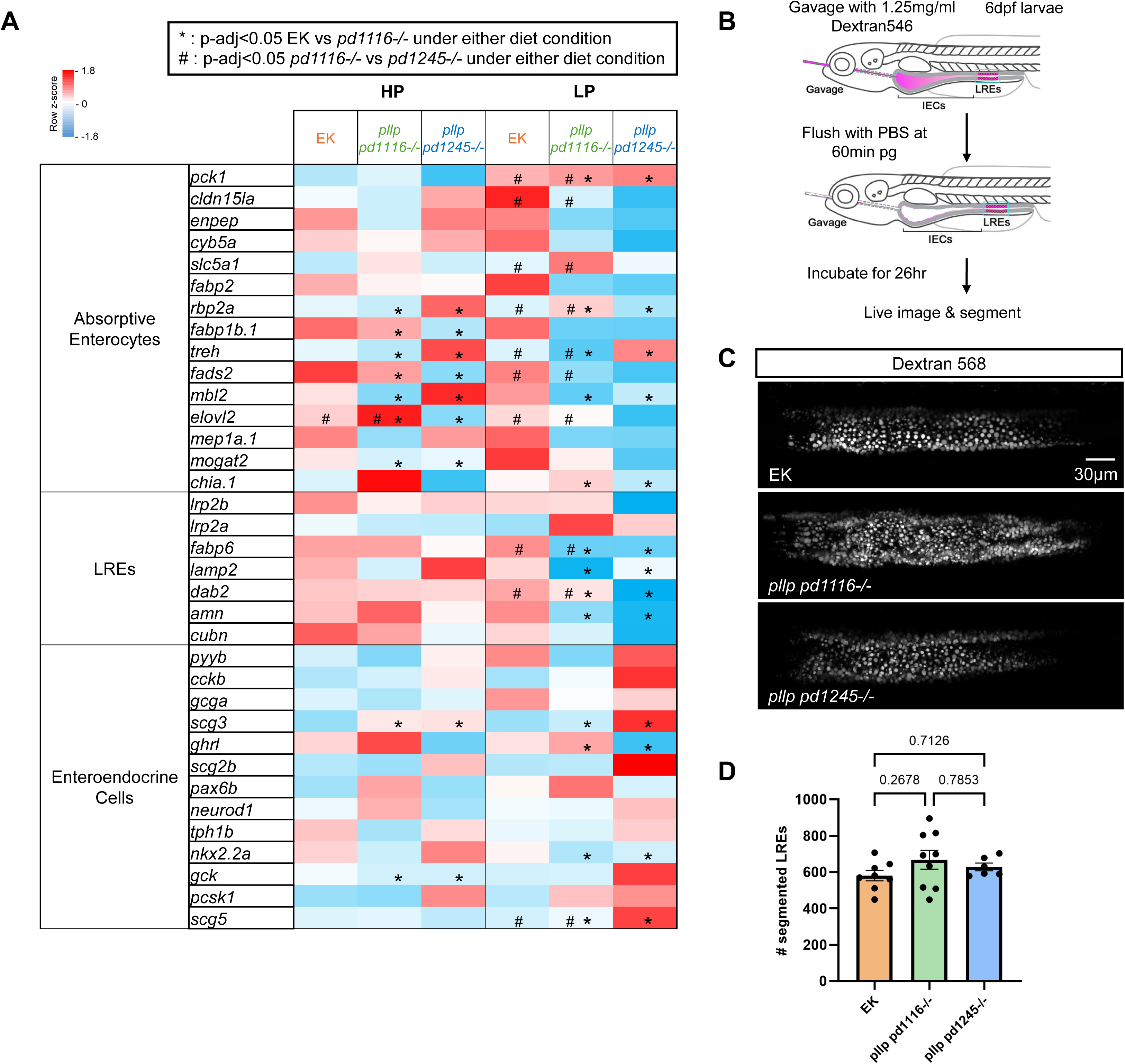
Normal LRE numbers and largely unaffected expression of genes characteristic for various intestinal cell populations in *pd1116-/-* and *pd1245-/-*. **(A)** Gene expression heatmap of genetic markers for the absorptive enterocytes, LREs, and enteroendocrine cells. **(B)** Experimental setup of active LRE quantification at 6dpf using dextran568 and using Ilastik object segmentation. **(C)** Representative images of the LRE regions showing dextran568-filled LRE lysosomal vacuoles. **(D)** Quantifications of active LRE cell numbers; n=8 for EK, n=9 for *pd1116-/*-, n=6 for *pd1245-/-*; one-way ANOVA.

**Figure S5.**
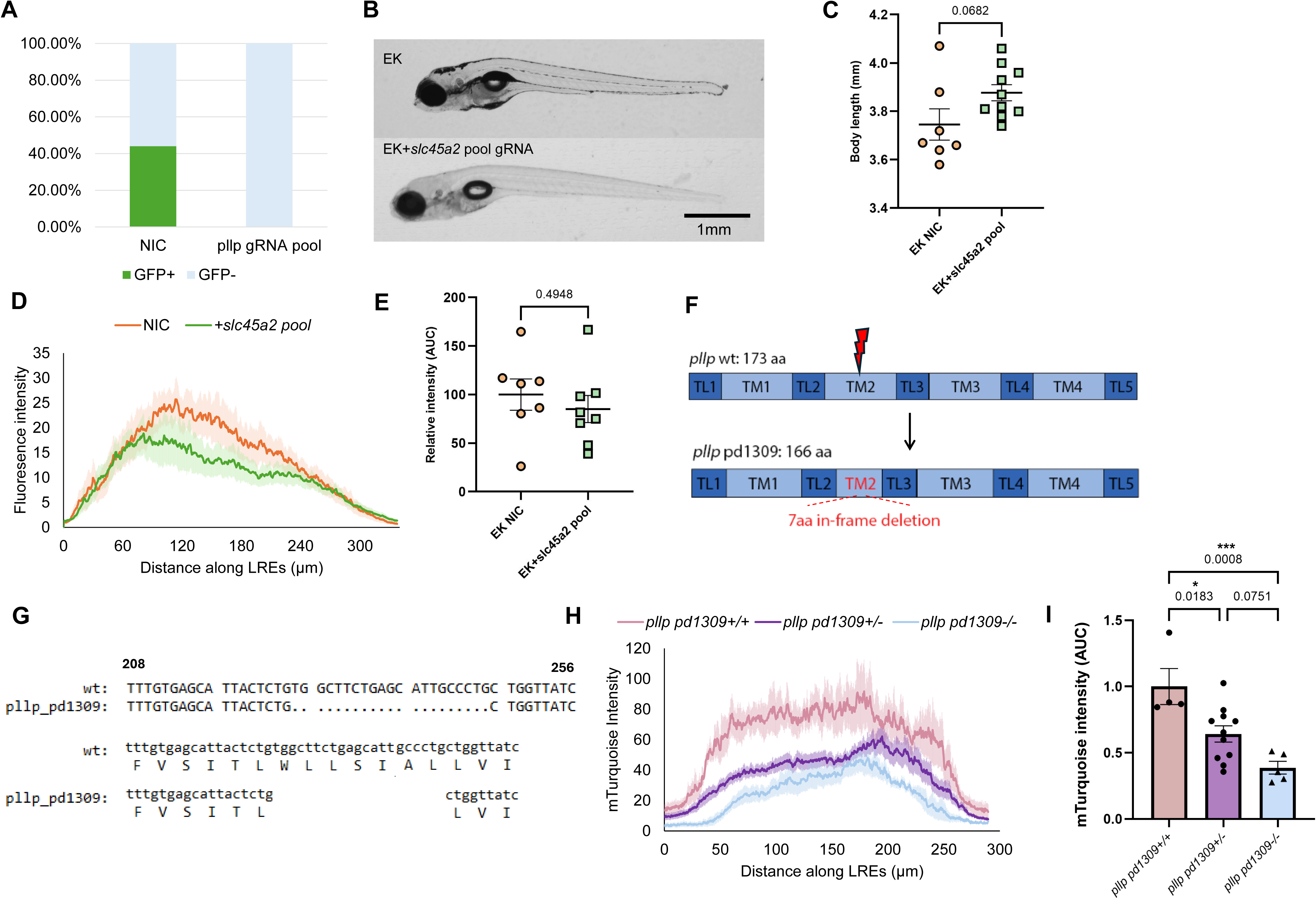
*pllp* CRISPR gRNA pool efficiency test, *slc45a2* CRISPR gRNA pool as a valid knock-out control and characterization of the *pd1309* allele. **(A)** Percentage of GFP+ *TgBAC(pllp^pd1114^-GFP)^+/-^ x EK* larvae either noninjected (NIC) or injected with the *pllp* CRISPR gRNA pool (EK+ *pllp* pool). Loss of GFP indicates that knockdown of *pllp* was efficient. **(B)** DIC images of EK and EK+*slc45a2* CRISPR gRNA pool; *slc45a2* knockout results in near complete loss of skin pigments. **(C)** Quantifications of body lengths of EK NIC and EK injected with *slc45a2* CRISPR gRNA pool. **(D, E)** Fluorescent profile (D) and quantifications (E) of Turquoise signal at 30min pg; n=7 for EK NIC; n=8 for EK+*slc45a2* pool; upaired t-test. **(F)** Diagram of the *pllp pd1309* allele generated by CRISPR-Cas9 single gRNA; editing results in a 21bp in-frame deletion within TM2. **(G)** DNA sequencing result (top) and predicted protein sequence (bottom) of the *pllp pd1309* allele compared to EK; numbers indicate bp downstream of the ATG start codon. **(H, I)** mTurquoise fluorescent signal profile (H) and quantifications (I) at 30min pg; n=4 *pd1309*+/+, n=11 for *pd1309*+/-, n=5 for *pd1309*-/-; data normalized to *pd1309*+/+; one-way ANOVA.

**Figure S6:**
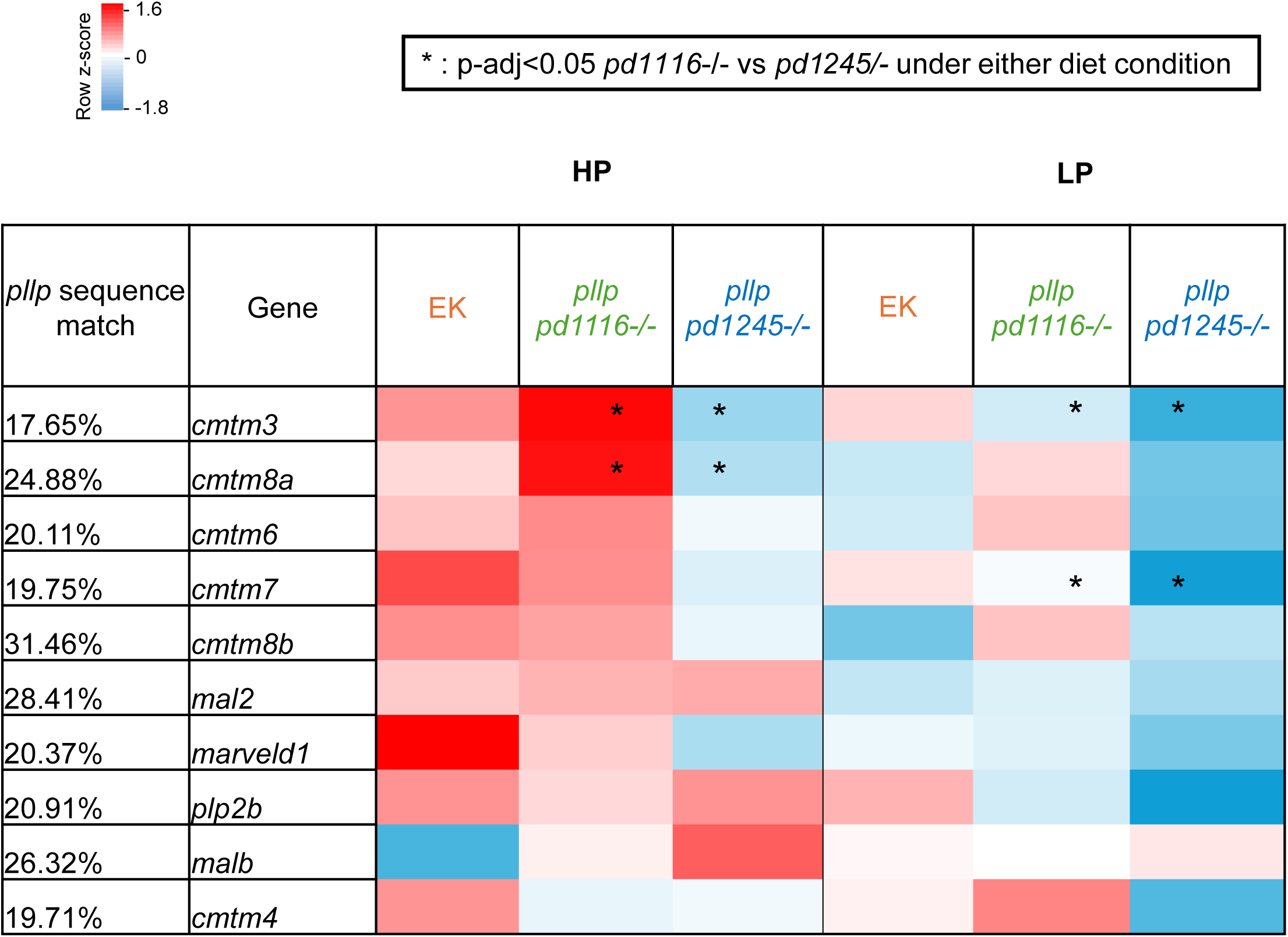
*pllp* paralog expression heatmap. *pllp* sequence match indicates the % of paralog sequences matching *pllp* sequence. No significant upregulation of any paralogs in pd1116-/- compared to EK. Sequence match data from ensembl.org.

## Notes

### Competing Interest Statement

The authors have declared no competing interest.

